# Profiling brain morphology for autism spectrum disorder with two cross-culture large-scale consortia

**DOI:** 10.1101/2025.02.24.639771

**Authors:** Xue-Ru Fan, Ye He, Yin-Shan Wang, Lei Li, the Lifespan Brain Chart Consortium (LBCC), the China Autism Brain Imaging Consortium (CABIC), Xujun Duan, Xi-Nian Zuo

## Abstract

We explore neurodevelopmental heterogeneity in Autism Spectrum Disorder (ASD) through normative modeling of cross-cultural cohorts. By leveraging large-scale datasets from Autism Brain Imaging Data Exchange (ABIDE) and China Autism Brain Imaging Consortium (CABIC), our model identifies two ASD subgroups with distinct brain morphological abnormalities: subgroup “L” is characterized by generally smaller brain region volumes and higher rates of abnormality, while subgroup “H” exhibits larger volumes with less pronounced deviations in specific areas. Key areas, such as the isthmus cingulate and transverse temporal gyrus, were identified as critical for subgroup differentiation and ASD trait correlations. In subgroup H, the regional volume of the isthmus cingulate cortex showed a direct correlation with individuals’ autistic mannerisms, potentially corresponding to its slower post-peak volumetric declines during development. These findings offer insights into the biological mechanisms underlying ASD and support the advancement of subgroup-driven precision clinical practices.

## INTRODUCTION

Autism Spectrum Disorder (ASD), or autism, is a lifelong neurodevelopmental condition characterized by impairments in social communication and the presence of repetitive, unusual sensorymotor behaviors^1,2^. The global prevalence of ASD is approximately 1% in children^3^, with different rates reported in specific regions, such as 2.8% in children aged 8 years in the United States^4^ and 0.7% in children aged 6 to 12 years in China^5^. Several factors, including improved survey methodologies, diagnostic practices, public awareness and access to services, have contributed to this increase in prevalence^3,6^. Over time, understanding of ASD has developed from a categorical diagnosis to a dimensional perspective of neurodiversity^7^. This change redefined autism as a spectrum that includes a wide range of characteristics within a unified framework^8^. However, the expanded diagnostic criteria may have too low thresholds for diagnosis^9^, and have introduced significant heterogeneity within ASD. This heterogeneity suggests that the term “spectrum” encompasses multiple subgroups with distinct etiological phenotypes^10,11^.

Recent neuroimaging studies increasingly highlight ASD’s biological heterogeneity as a key barrier to identifying consistent brain-behavior relationships. Traditional case-control neuroimaging studies, which assume ASD and typical developing groups as homogeneous entities, often fail to account for subgroup differences^12^. This has led to inconsistent findings across brain regions and participants, further complicating efforts to identify reliable brain-behavior biomarkers. The decreasing effect sizes in group comparison studies over time^13^ highlight the difficulty of constructing mechanistic models of ASD. This heterogeneity challenge extends to intervention research, where differential treatment responses across ASD subgroups^14,15^ suggest that precision approaches accounting for neurobiological variation could enhance outcomes. Data-driven clustering techniques have emerged as a promising approach to address this complexity by leveraging cognitive and behavioral profiles alongside patterns of brain morphology and function. Various clustering methods^16–20^ have been used to identify meaningful subgroups, in order to uncover the neurobiological mechanisms of ASD and provide insights into subgroup-specific traits^8,21^. While despite decades of effort, inconsistencies in sample selection, classification methods, and feature selection between studies remain significant challenges^21^. These issues have limited the ability of clustering techniques to consistently identify robust and replicable subgroups.

Inconsistencies in ASD studies may be due to altered neurodevelopmental adaptations^22^ and participant selection biases^23^. These limitations highlight the need to group variables with strong biological relevance for the ASD subgroups^24^. At the same time, individual differences in sex, intelligence, medication use, and comorbidities^25,26^, along with methodological variability^27^ in data acquisition and analysis, complicate the interpretation of the results of neuroimaging studies. In response to these challenges, normative modeling^28^ techniques offer a promising framework to capture the full range of neurobiological variation^12^. This approach enables standardized assessments by comparing an individuals brain morphology to population-based references^29^. In previous ASD studies, references were often built on internal or public datasets^18,20,30,31^, but the relatively small sample size limits the generalizability and reliability of the results. Recently, the Lifespan Brain Chart Consortium (LBCC, https://github.com/brainchart/lifespan) provided neurodevelopmental variations throughout life from the most inclusive data available^32^, providing a robust framework for us to achieve percentile-based comparisons of ASD with considerable statistical power.

This study adjusted the LBCC normative models in two large-scale cross-cultural ASD consortia: the China Autism Brain Imaging Consortium (CABIC)^33^ and the Autism Brain Imaging Data Exchange (ABIDE)^34,35^. Focusing on childhood development, we used the largest-scale brain charts^32^ to explore neurodevelopmental heterogeneity in ASD before adolescence. We classified ASD into subgroups using spectral clustering based on Out-of-Sample (OoS) centile scores, a biological measure which quantified brain morphological deviations of ASD from normative growth. A Support Vector Machine (SVM) with Recursive Feature Elimination and Cross-Validation (RFECV) identified key brain regions driving subgroup classification. The ABIDE-based classifier was then applied to CABIC to identify similar subgroups. Brain-behavior analyses were performed separately for each dataset to identify reproducible subgroup-specific correlations. From a perspective of brain morphology, this work disentangles the mechanisms of mixed neurosubtypes and provides new insight into the biological complexity of ASD.

## RESULTS

### Brain Morphological Profiles Reveal Two Distinct Autism Subgroups

We first analyzed the ABIDE dataset. The distributions of the OoS centile scores across 34 bilateral Desikan-Killiany^36^ cortical volumes ranged from 0 to 1 (Figure S1a in the Supplementary Information). OoS scores estimate the morphological variations of each participant in 34 brain regions compared to the median regional volumes of the population (50th centile, equivalent to a 0.5 OoS score). Relative to 0.5, individuals with more atypical phenotypes have more extreme scores. A Shapiro-Wilk test on the 34 OoS scores indicated that none followed a normal distribution (Table 1, column 2). Using spectral clustering on the OoS scores, we identified two distinct clusters (ASD subgroups) with the highest silhouette coefficient among cluster solutions ranging from two to ten (Figure 1a). All *p*-values for between-subgroup comparisons (two-sample *t*-tests for the paracentral and pars orbitalis regions, and rank-sum tests for all other regions) were *<* 0.001). One subgroup, referred to as “L”, exhibited generally *Lower* OoS scores, while the other referred to as “H”, displayed *Higher* scores (Table 1, columns 5 and 7).

**Table 1:** Group comparisons of regions morphology between overall and two ASD subgroups.

| Region | OoS Scores |  |  |  |  |  | AbP<sup>1</sup>(%) |  |
| --- | --- | --- | --- | --- | --- | --- | --- | --- |
|  | All |  | L |  | H |  | L | H |
|  | *p* | Median | *p* | Median | *p* | Median |  |  |
| Superior frontal | 0.01 | 0.48 | 0.02 | 0.28 | 0.37 | 0.62 | 5.56 | 2.70 |
| Rostral middle frontal | 0.01 | 0.46 | 0.02 | 0.25 | 0.32 | 0.60 | 7.94 | 0.68 |
| Caudal middle frontal | 0.00 | 0.42 | 0.00 | 0.26 | 0.05 | 0.60 | 6.35 | 2.70 |
| Pars opercularis | 0.00 | 0.47 | 0.00 | 0.23 | 0.07 | 0.61 | 6.35 | 4.73 |
| Pars triangularis | 0.00 | 0.45 | 0.02 | 0.28 | 0.02 | 0.60 | 6.35 | 1.35 |
| Pars orbitalis | 0.01 | 0.46 | 0.06 | 0.33 | 0.11 | 0.56 | 5.56 | 0.68 |
| Frontal pole | 0.00 | 0.48 | 0.02 | 0.34 | 0.03 | 0.60 | 11.90 | 2.03 |
| Lateral orbital frontal | 0.00 | 0.46 | 0.01 | 0.26 | 0.06 | 0.63 | 7.94 | 2.03 |
| Medial orbital frontal | 0.00 | 0.42 | 0.00 | 0.26 | 0.04 | 0.61 | 10.32 | 2.70 |
| Rostral anterior cingulate | 0.00 | 0.45 | 0.00 | 0.25 | 0.04 | 0.65 | 4.76 | 2.70 |
| Caudal anterior cingulate | 0.00 | 0.47 | 0.01 | 0.30 | 0.00 | 0.68 | 3.97 | 6.76 |
| Precentral | 0.00 | 0.45 | 0.01 | 0.26 | 0.17 | 0.60 | 6.35 | 3.38 |
| Paracentral | 0.01 | 0.52 | 0.05 | 0.34 | 0.10 | 0.62 | 4.76 | 5.41 |
| Postcentral | 0.00 | 0.48 | 0.01 | 0.29 | 0.05 | 0.64 | 7.14 | 4.05 |
| Supramarginal | 0.00 | 0.49 | 0.01 | 0.31 | 0.04 | 0.65 | 7.94 | 3.38 |
| Posterior cingulate | 0.00 | 0.46 | 0.02 | 0.29 | 0.04 | 0.64 | 7.94 | 5.41 |
| Isthmus cingulate | 0.00 | 0.47 | 0.00 | 0.27 | 0.06 | 0.64 | 3.97 | 4.05 |
| Precuneus | 0.00 | 0.52 | 0.04 | 0.29 | 0.02 | 0.70 | 5.56 | 4.73 |
| Superior parietal | 0.00 | 0.44 | 0.00 | 0.27 | 0.08 | 0.60 | 7.94 | 2.70 |
| Inferior parietal | 0.00 | 0.45 | 0.07 | 0.30 | 0.02 | 0.63 | 7.94 | 2.03 |
| Transverse temporal | 0.00 | 0.52 | 0.02 | 0.32 | 0.03 | 0.68 | 7.94 | 6.08 |
| Banks superior temporal | 0.00 | 0.44 | 0.00 | 0.26 | 0.02 | 0.59 | 3.97 | 5.41 |
| Superior temporal | 0.00 | 0.49 | 0.00 | 0.26 | 0.04 | 0.67 | 6.35 | 4.05 |
| Middle temporal | 0.00 | 0.44 | 0.02 | 0.24 | 0.06 | 0.61 | 17.46 | 2.70 |
| Inferior temporal | 0.00 | 0.49 | 0.05 | 0.30 | 0.04 | 0.62 | 11.90 | 0.68 |
| Fusiform | 0.00 | 0.45 | 0.01 | 0.26 | 0.09 | 0.59 | 10.32 | 1.35 |
| Parahippocampal | 0.00 | 0.49 | 0.02 | 0.33 | 0.01 | 0.63 | 3.17 | 2.70 |
| Entorhinal | 0.01 | 0.51 | 0.05 | 0.40 | 0.04 | 0.62 | 6.35 | 0.68 |
| Temporal pole | 0.00 | 0.44 | 0.03 | 0.36 | 0.02 | 0.58 | 10.32 | 1.35 |
| Lateral occipital | 0.00 | 0.46 | 0.01 | 0.28 | 0.02 | 0.63 | 9.52 | 2.70 |
| Lingual | 0.00 | 0.40 | 0.00 | 0.24 | 0.01 | 0.58 | 7.94 | 2.70 |
| Pericalcarine | 0.00 | 0.36 | 0.00 | 0.23 | 0.01 | 0.53 | 11.90 | 2.70 |
| Cuneus | 0.00 | 0.46 | 0.01 | 0.25 | 0.03 | 0.55 | 6.35 | 1.35 |
| Insula | 0.00 | 0.53 | 0.02 | 0.31 | 0.02 | 0.69 | 3.17 | 6.08 |

<sup>1</sup>AbP: Prevalence of abnormalities.

**Figure 1:**
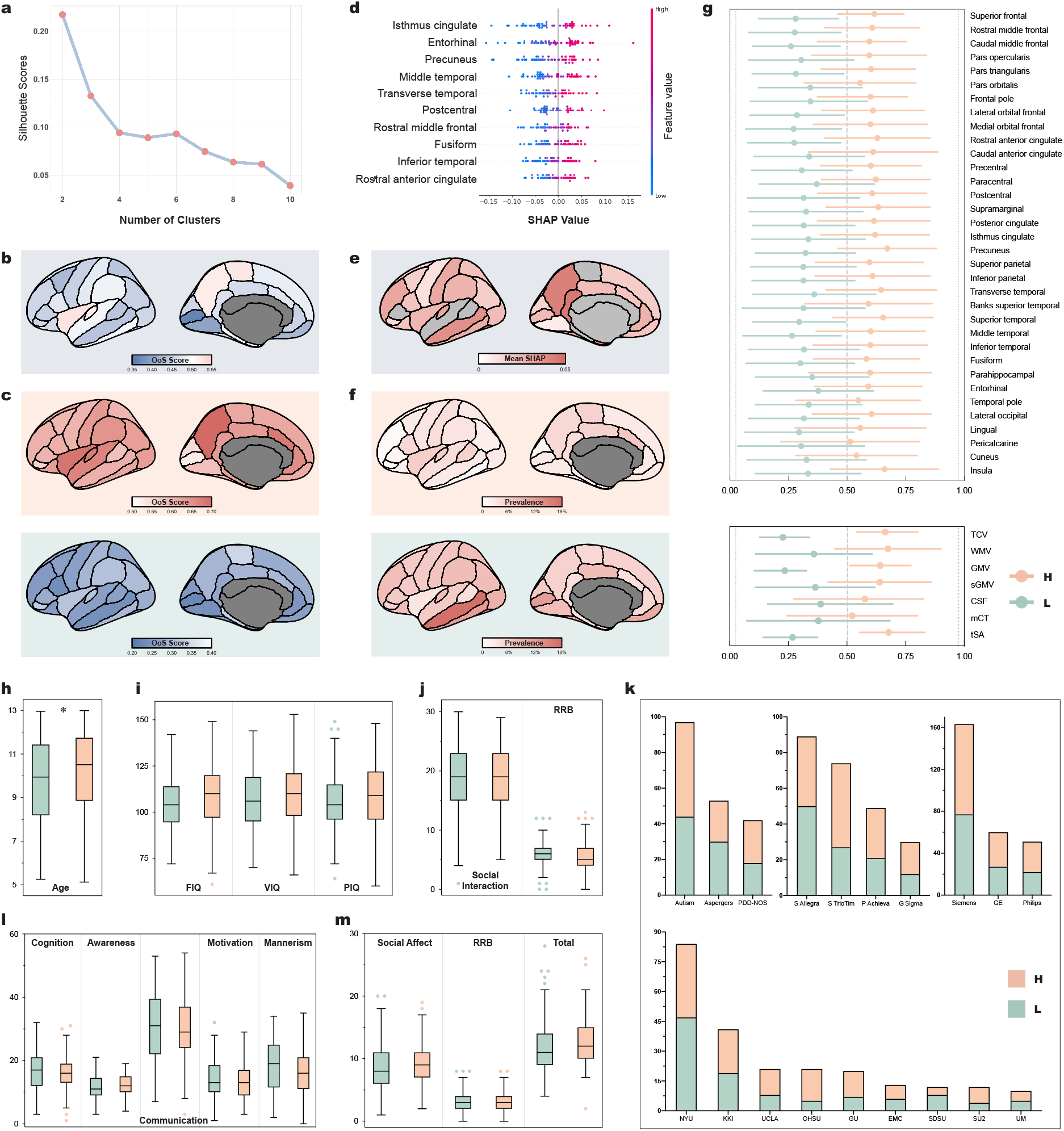
Clustering-derived subgroups reveal distinct brain morphological patterns in ASD from ABIDE dataset. **a**, Silhouette scores across different clustering solutions (2 to 10 clusters). The highest silhouette score indicates two distinct subgroups. **b**, Median OoS scores of 34 cortical regions before clustering. **c**, Median OoS scores of ASD subgroup H (top) and L (bottom). **d**, SHAP summary plot displaying the top 10 brain regions with the highest contributions to the SVM model’s predictions. **e**, The mean SHAP values across the 29 selected cortical regions contributed to the classification. **f**, Prevalence maps depicting the proportion of participants with extreme (2.5% for subgroup L, 97.5% for subgroup H) structural anomalies. **g**, OoS scores across 34 cortical regions (top) and global measures (bottom) for subgroup H (orange) and L (green). TCV, total cortical volume; WMV, total white matter volume; GMV, total cortical gray matter volume; sGMV, total subcortical gray matter volume; CSF, total ventricular cerebrospinal fluid volume; mCT, mean cortical thickness; tSA, total surface area; Dots indicate the mean value of OoS scores, bars indicate the standard deviation. See Supplementary Information Figure S1b and Figure S1d for detailed density plots. Distribution of participant ages (**h**); IQ scales (**i**); ADI-R (**j**); PDD category (**k**, top left), MRI scanner model (**k**, top middle), manufacturer (**k**, top right), and data collection site (**k**, bottom); SRS (**l**); and ADOS (**m**) across the two subgroups, with subgroup L participants being significantly younger than those in subgroup H (*p* = 0.02). Note, the left hemispheres are plotted in **b, c, e**, and **f** just for visualization purposes. For plots **h, i, j, l**, and **m**, the center line shows the median; the box limits represent the 25th and 75th percentiles; the whiskers show the minimum and maximum values; and the dots represent potential outliers.

Certain regions of the brain showed significant differences in their OoS score distributions between the two subgroups (Figure 1g; Figure S1b in the Supplementary Information. For CABIC results, please see Figure S2a and S2b). Subgroup H exhibited greater variability in the occipital lobe, particularly in the lingual gyrus, pericalcarine cortex, and cuneus. In contrast, subgroup L showed more pronounced reductions in these regions compared to the normative benchmark (0.5). Other regions, such as the pars opercularis and superior temporal gyrus, also exhibited subgroup-specific changes. The results of the Shapiro-Wilk test for OoS scores (Table 1, columns 4 and 6) indicated that more brain regions in subgroup H followed a normal distribution compared to subgroup L. Broader variability was observed in regions such as the entorhinal cortex in subgroup L and the pericalcarine in subgroup H, high-lighting further subgroup-specific clustering.

### Distinct Structural and Age Differences Between Subgroups

The median OoS scores before clustering (Figure 1b; Table 1, column 3) reveal a mixed pattern of deviations across regions. Most regions exhibit reduced volumes, particularly in the occipital lobe. However, some regions, such as the precuneus, paracentral lobule, transverse temporal gyrus, and insula, show slightly higher OoS scores. This combined pattern reflects overlapping structural deviations from both subgroup H (Figure 1c, top) and L (Figure 1c, bottom), obscuring the distinct morphological differences that become evident after clustering. For instance, although subgroup L shows smaller volumes in the insula and precuneus, the larger volumes in subgroup H predominantly drive the combined pattern in the whole population. In contrast, smaller volumes of the lingual gyrus and the middle temporal gyrus in subgroup L have a stronger influence on the overall result. Some regions, such as the pericalcarine cortex, are almost entirely influenced by subgroup L. Meanwhile, regions like the inferior and superior temporal gyrus exhibit volumes close to the normative average because of the contrasting characteristics of both subgroups effectively canceling each other out.

To identify the brain regions most associated with morphological abnormalities, we calculated the prevalence of abnormalities for each region in the two subgroups. The prevalence represents the percentage of participants with OoS scores below 0.025 (2.5% centile) for subgroup L, while above 0.975 (97.5% centile) for subgroup H. Subgroup L (Figure 1f, bottom; Table 1, column 8) exhibited the highest prevalence of abnormalities in regions such as the middle and inferior temporal gyrus, pericalcarine cortex, frontal pole, and medial orbital frontal gyrus. In contrast, subgroup H (Figure 1f, top; Table 1, column 9) generally exhibited less severe abnormalities, with a concentration in the insula, transverse temporal gyrus, and caudal anterior cingulate. Participants in subgroup L were significantly younger than those in subgroup H (Figure 1h; *p* = 0.02, Table S1 in Supplementary Information). However, no significant differences were observed in the category PDD (*p* = 0.32), the MRI scanner model (*p* = 0.07) and the manufacturer (*p* = 0.86), the data collection site (*p* = 0.16) (Figure 1k), IQ (Figure 1i, see Table S1 in Supplementary Information for details), or scores from ADI-R (Figure 1j), SRS (Figure 1l), and ADOS-2 (Figure 1m). The corresponding results for CABIC are shown in Supplementary Information Figure S3.

### Machine Learning Reveals Subgroup Morphological Features

The optimized SVM model for the identification of subgroups demonstrated robust predictive performance. It used a polynomial kernel with a regularization parameter *C* = 0.1 and a kernel coefficient *γ* = 1. The model achieved a high classification accuracy (0.95, *p <* 0.001) and a F1 score (0.94, *p <* 0.001) in five cross-validation folds. Figure 1d shows the SHapley Additive exPlanations (SHAP) summary plot for the top 10 selected features. Figure 1e and Table S2 in Supplementary Information summarize the mean SHAP values for all 29 selected regions. SHAP values^37^ quantify the contribution of individual brain regions to the predictions of the model. Among the regions, the isthmus cingulate, entorhinal cortex, precuneus, and middle temporal gyrus emerged as the most predictive features. Higher SHAP values for these regions were strongly associated with subgroup H, while lower values corresponded to subgroup L. These findings indicate different volumetric patterns in the cingulate, temporal, and parietal areas, effectively distinguishing the two subgroups.

### Distinct Structural Covariance Patterns Across Subgroups

Structural covariance analysis on OoS scores revealed significant correlations between brain regions, with distinct patterns observed across ASD subgroups and control group (Figure 2c). These patterns were reproduced in both the ABIDE and CABIC datasets. In subgroup H, enhanced covariation was identified between the isthmus cingulate and the caudal middle frontal region (ABIDE: *z* = 2.88, *p* = 0.00; CABIC: *z* = 2.25, *p* = 0.02). In subgroup L, the parahippocampal gyrus showed reduced covariation with the entorhinal cortex (ABIDE: *z* = −2.07, *p* = 0.04; CABIC: *z* = −3.25, *p* = 0.00). Similarly, reduced covariation was observed between the posterior cingulate and the entorhinal cortex (ABIDE: *z* = −2.12, *p* = 0.03; CABIC: *z* = −2.64, *p* = 0.01) and between the insula and the entorhinal cortex (ABIDE: *z* = −3.30, *p* = 0.00; CABIC: *z* = −2.18, *p* = 0.03). An increased covariation was observed in subgroup L between the parahippocampal gyrus and the lateral occipital cortex (ABIDE: *z* = 1.97, *p* = 0.05; CABIC: *z* = 2.04, *p* = 0.04).

**Figure 2:**
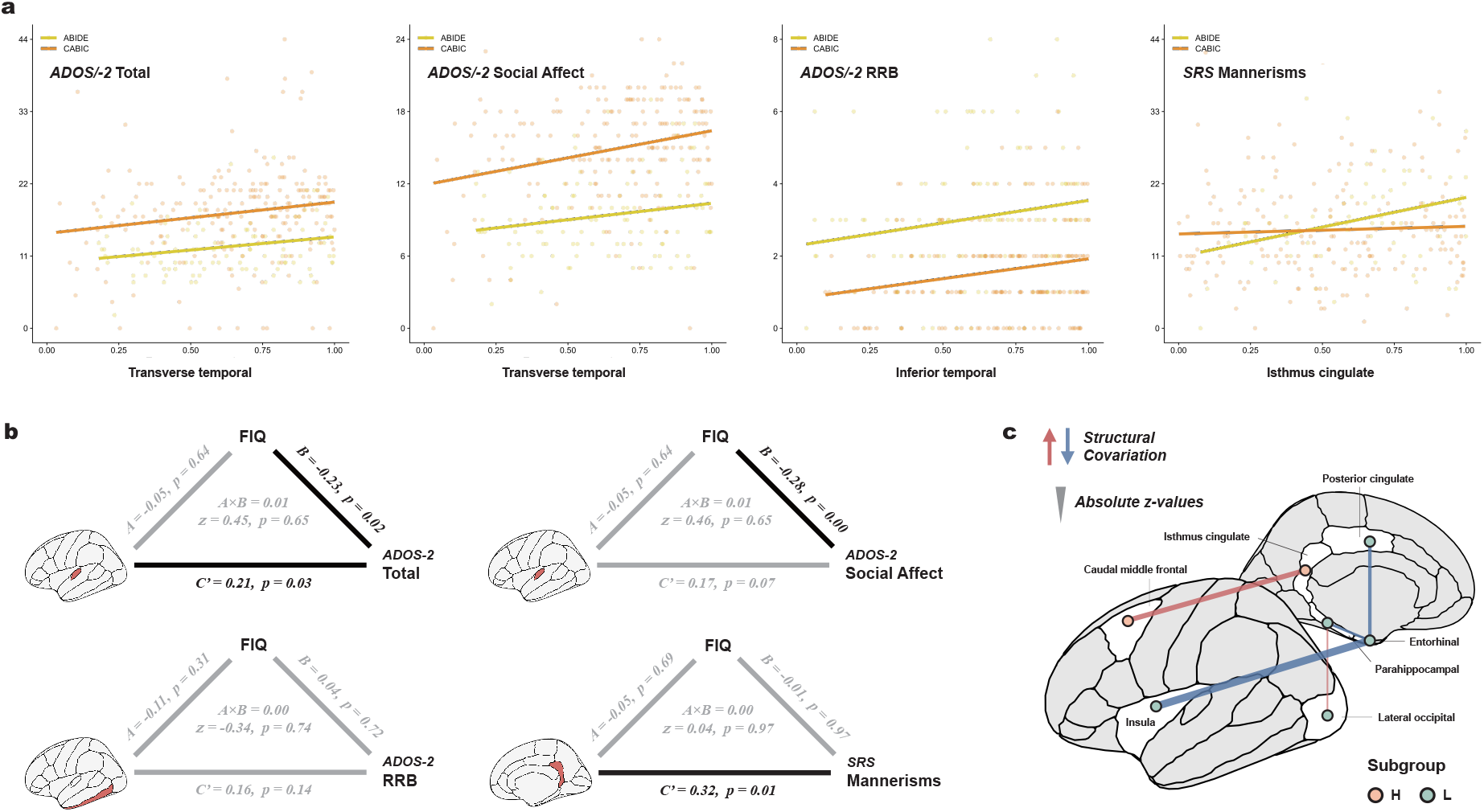
Brain-behavior correlations and structural covariations of Out-of-Sample scores. **a**, Reproducible correlations between brain region volumes, measured as Out-of-Sample (OoS) scores, and clinical measures across ABIDE and CABIC datasets (subgroup H only). **b**, Mediation models for significant brain-behavior associations identified in subgroup H (ABIDE cohort only). Black solid arrows represent significant effects, while gray arrows indicate non-significant ones. Top: transverse temporal; bottom left: inferior temporal; bottom right: isthmus cingulate. **c**, Significant structural covariations across ABIDE and CABIC datasets. Positive *z*-values (red lines) indicate stronger covariance in ASD participants compared to controls, while negative *z*-values (blue lines) reflect weaker covariance.

### Brain-Behavior Correlations in Subgroup H

We examined correlations between cortical region volumes (OoS scores) and clinical measures in the ABIDE and CABIC datasets, identifying significant associations that were consistent between both datasets. All significant correlations were found within subgroup H. The transverse temporal gyrus showed positive correlations with ADOS (ADOS-2 in ABIDE and ADOS in CABIC) Total and Social Affect scores (Figure 2a). In the CABIC dataset, moderate correlations were observed for Total (*r*(198) = 0.17, *p* = 0.02) and Social Affect (*r*(191) = 0.21, *p* = 0.00) score. Stronger correlations were identified in the ABIDE dataset for Total (*r*(93) = 0.24, *p* = 0.01) and Social Affect (*r*(93) = 0.20, *p* = 0.04) score. The inferior temporal gyrus was positively correlated with RRB (restricted interests and repetitive behaviors), with a slightly stronger correlation in ABIDE (*r*(94) = 0.19, *p* = 0.05) compared to CABIC (*r*(189) = 0.17, *p* = 0.02). The volumetric centile of the isthmus cingulate is correlated with SRS Autistic Mannerisms, showing a higher correlation in ABIDE (*r*(58) = 0.33, *p* = 0.01) than in CABIC (*r*(192) = 0.14, *p* = 0.04). No significant correlations were identified between structural covariance patterns and cognitive behavior outcomes.

As a common comorbidity of ASD, intellectual disability is closely associated with atypical brain morphological developmental patterns that accompany individual differences in intellectual functioning. To examine whether Full-Scale Intelligence Quotient (FIQ) mediates the relationships between brain morphology and cognitive behaviors within clusters, mediation analyses were conducted on the significant correlations we found in subgroup H. Results revealed direct effects for specific cortical regions (Figure 2b). The transverse temporal gyrus showed a direct effect on ADOS-2 Total (*C*^*′*^ = 0.21, *p* = 0.03), while the isthmus cingulate exhibited a direct effect on SRS Autistic Mannerisms (*C*^*′*^ = 0.32, *p* = 0.01). As our analysis relied on cross-sectional data, the observed brainbehavior relationships represent correlations rather than causal effects. Our mediation hypotheses regarding developmental brainbehavior causality require validation through longitudinal studies. While no mediation effects through FIQ were observed, both from brain → cognitive behaviors or in reverse (Figure S4). Neither direct nor indirect effects were significant for correlations to ADOS2 RRB or Social Affect.

## DISCUSSION

This study used the ABIDE and CABIC datasets to discover distinct brain morphological subgroups of ASD within a young male cohort. We identified two subgroups characterized by significant differences in OoS scores in 34 cortical regions. Machine learning models showed a high predictive accuracy in distinguishing these subgroups based on their brain morphology. Furthermore, structural covariance and brain behavior correlation analysis illustrated different patterns of morphological relationships and their associations with clinical measures, especially in subgroup H. The consistent correlations across both the ABIDE and CABIC datasets highlight the robustness and reproducibility of our results across diverse cohorts. We will now discuss these findings from several neuro-biological and neurodevelopmental perspectives to enhance our understanding of the distinct morphological patterns identified.

### Two Subgroups Suggest Different Neurobiological Basis

Studies have reported extremely large or small head circumferences in individuals with ASD, possibly related to different subgroups. Early brain overgrowth is one of the most consistent findings in ASD research^38^. Abnormal increases in brain size during early development suggest that atypical cell proliferation significantly contributes to ASD symptoms^39^. Ultrasound studies have detected head enlargement during the second trimester in individuals later diagnosed with ASD^40^. iPSC-based studies reveal enlarged embryonic stage brain cortical organoids (BCOs) in babies with ASD, with larger BCOs associated with more severe social symptoms^41^. MRI scans of these individuals show overgrowth in primary auditory and somatosensory cortices, while undergrowth in the visual cortex^41^. This neuronal overproduction, caused by the acceleration of the cell cycle, results in impaired differentiation and ultimately disrupts neural functions^42^.

Smaller brain volumes are also frequently reported in ASD. A meta-analysis^43^ identified volumetric reductions in a large cluster of regions, including the parahippocampal gyrus and entorhinal cortex. Higher levels of autistic traits have been associated with a smaller total cortical volume (TCV), a lower cortical thickness, a smaller surface area and a lower gyrification^44,45^. One potential explanation for this global brain underdevelopment is impairment caused by insufficient blood circulation and oxygen saturation^46^. For example, children with complex congenital heart disease (CHD) also demonstrate cognitive difficulties similar to those observed in ASD and have an increased likelihood of developing ASD^47^. Similarly, head enlargement also coincide with greater increases in height often^45^. Therefore, brain overgrowth could be part of the broader physical growth dysregulation^48^ too.

These evidences provide the distinct neurobiological basis for the subgroups identified in our study. The different underlying physiological mechanisms may explain the inconsistent effectiveness of commonly used pharmacological treatments. For example, unsupervised data-driven cluster analysis on ASD children revealed an optimum of two intranasal oxytocin intervention-sensitive subtypes^49^. Our discovery of two distinct ASD subtypes with divergent brain morphologies further supports the necessity for subtypespecific therapeutic strategies. Future research focusing on targeted pharmacological interventions for individual subtypes will not only explain how these treatments modulate brain function and ultimately translate into clinical benefits, but also advance personalized medicine in ASD. Specifically, by mapping neurobiological heterogeneity to differential therapeutic outcomes, this approach could reconcile previous inconsistencies in treatment efficacy while optimizing intervention protocols for mechanistically defined patient subgroups.

### Hypothesis of Abnormal Brain Morphology Specific Functional Impairment Pathways

Abnormal neural migration and cortical laminar organization in ASD^50^ suggest that structural abnormalities are unlikely to be localized. Postmortem studies have identified focal laminar disorganization and mis-migrated neurons in the prefrontal and temporal cortex, particularly in layers 2, 3, and 4^50^. Layers 2 and 3 support information exchange between cortical regions, while layer 4 receives sensory inputs^51^. Disruptions in these layers can cause profound miscommunication between the sensory and higher-order cognitive regions. Key regions in our clustering analysis (such as the precuneus, isthmus cingulate, and entorhinal cortex; Figure 3a, *a*∼*c*) are primarily involved in higher-order association networks (Figure 3a). As an idea for exploring hierarchical information flow of functional connectivity, Stepwise Functional Connectivity (SFC) analysis was introduced to disentangle brain functional networks, and has discovered the complex connectivity transitions from primary sensation to higher-order association regions of the brain^52,53^. These existing information flow suggest different pathways of impairment as the basis for the heterogeneity of ASD. Inconsistencies in information flow reported by previous ASD studies^21,54^ may arise from two distinct structural impairment patterns identified in our study.

**Figure 3:**
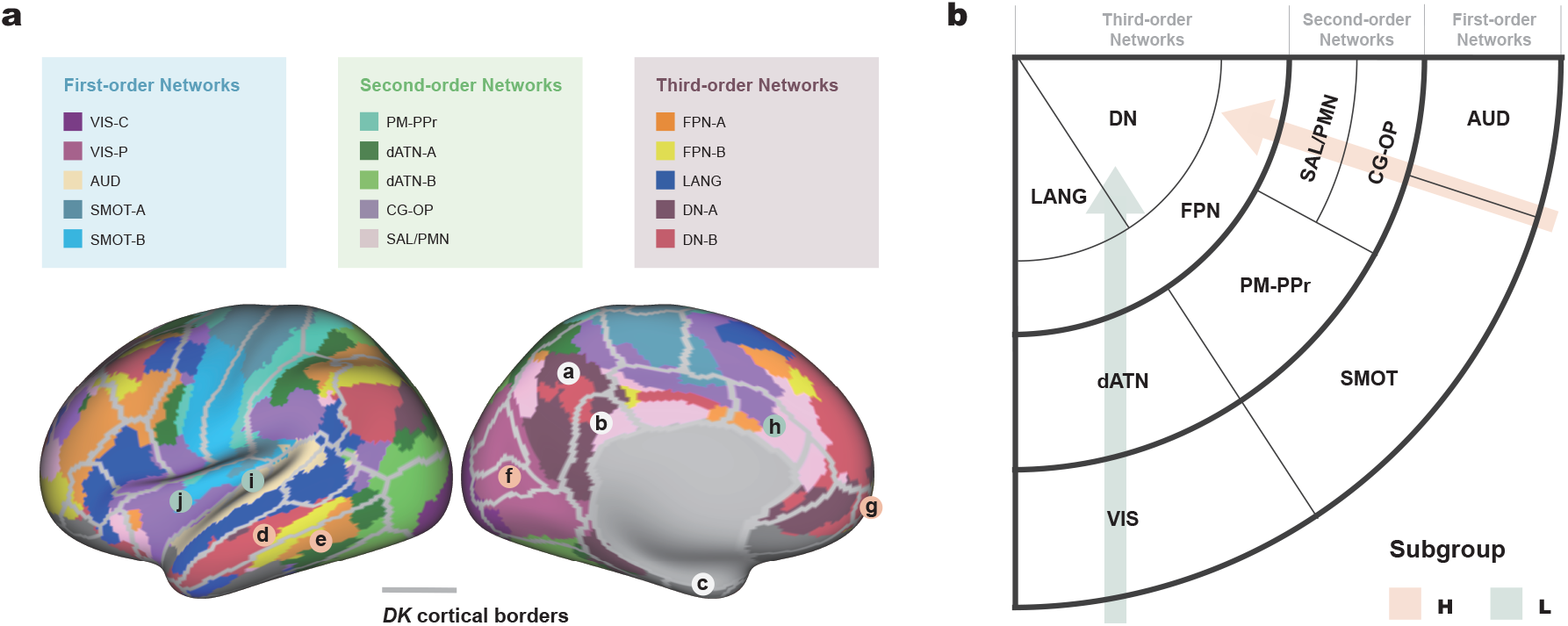
The theoretical brain functional network impairment model of two ASD subgroups. **a**, A brain map combining the 34 Desikan regions^36^ with the latest 15 large-scale brain functional networks estimated from individuals^57^. *a*, precuneus; *b*, isthmus cingulate; *c*, entorhinal; *d*, middle temporal; *e*, inferior temporal gyrus; *f*, pericalcarine; *g*, frontal pole; *h*, caudal anterior cingulate; *i*, transverse temporal; and *j*, insula. **b**, A schematic of the potential spatial distributions of distinct functional impairment networks inferred from 34 cortical regions with high abnormal prevalence for two subgroups. VIS, Visual (C, Central; P, Peripheral); SMOT, Somatomotor; AUD, Auditory; dATN, Dorsal Attention; PM-PPr, Premotor-Posterior Parietal Rostral; CG-OP, Cingulo-Opercular; SAL/PMN, Salience/Parietal Memory; FPN, Frontoparietal; LANG, Language; DN, Default networks.

The regional volume abnormalities identified in our study involve distinct functional brain regions in the two subgroups. Subgroup L showed a notable prevalence in the middle temporal gyrus, inferior temporal gyrus, pericalcarine cortex and frontal pole (Figure 3a, *d*∼*g*), while subgroup H in the caudal anterior cingulate, transverse temporal cortex, and insula (Figure 3a, *h*∼*j*). Drawing on the spatial distributions^55^ of 7 large-scale functional networks^56^, we constructed a schematic (Figure 3b) of the latest 15 large-scale functional networks of the brain estimated from individuals^57^ (Figure 3a). This perspective revealed different pathways of functional impairment in the regions with a high prevalence of abnormalities for each subgroup. Subgroup L exhibited an atypical morphology mainly affecting Visual – Dorsal Attention – Frontoparietal – Language and Default networks (Figure 3b, green path), while subgroup H affected the Auditory and Somatomotor – Cingulo-Opercular and Salience/Parietal Memory – Frontoparietal networks (Figure 3b, orange path), indicating disturbances in peculiar unimodal sensory integration and multimodal cortical functions. These networks are organized by three-order hierarchy that are agree well with myelination reference maps^57^, showing that the cerebral cortex develops sequentially, radiating outward from motor and sensory cortex^58^.

This phenomenon may hint at the presence of distinct abnormal brain functional circuits from first-to third-order networks in the subgroups. For example, in adults without ASD^59^, increased gray matter volume in regions of the somatomotor network was associated with greater attention to detail, while changes in regions of the visual network were associated with poor imagination. These findings suggest potential cognitive patterns in two subgroups of ASD that have yet to be discovered. At the same time, these two subgroups we found may respond differently to biologically targeted therapies, emphasizing the need for more personalized approaches. For example, functional connectivity-guided continuous theta-burst stimulation (cTBS) on individual’s dorsolateral prefrontal cortex target with the strongest connectivity to amygdala significantly improve social communication skills in minimally verbal ASD children^60^. As reduction in brain volume and improvements in behavior outcomes, this ASD subtype likely fit the characteristics of subgroup H in our study. Targeted modulation of subgroup specific impairment pathway may gain better therapeutic effects, therefore further refine treatment protocols for better translational outcomes. In this case, if observed social improvements stem from rebalancing the subgroup H pathway, future interventions could achieve enhanced treatment efficacy by employing personalized neuromodulation protocols that precisely target the most morphologically burdened brain regions within this circuitry.

In typically developing children, the principal functional gradient^55^ of the cortical organization reflects differentiation within unimodal sensory areas, progressing from the somatomotor and auditory cortex to the visual cortex. The transition from childhood to adolescence extends this organization to higher-order association cortices^61^. Brain maturation appears to involve a shift from local to distributed network architecture. The distinct morphological abnormalities in these two ASD subgroups, observed before this transition begins, raise a critical question: How do these “morphological foundations” disrupt their development from childhood to adolescence? These structural impairment pathways in childhood are likely to contribute further to heterogeneity in ASD during and after adolescence.

### Understanding the Heterogeneity from a Developmental Perspective

Different network-based vulnerabilities suggest that interventions targeting specific impairments within each subgroup could be more effective. Interestingly, we found no significant differences in cognitive or behavioral measures between the subgroups. Although there distinct patterns of structural covariation between subgroups, but no significant correlations with autism-related behaviors. Structural covariance reflects long-term developmental changes. Reduced structural covariance may indicate disruptions in shared developmental pathways, while enhanced covariance could result from compensatory adaptations in response to atypical neurodevelopmental trajectories in ASD. This implies that morphological deficiencies may activate compensatory mechanisms during development, leading to similar cognitive and behavioral outcomes.

These neurodevelopmental compensation mechanisms raise questions about the stability of subgroup morphology distinctions across different developmental stages. To validate the robustness of our ASD clusters across different development stage, we implemented sliding 4-year age windows (with a predefined threshold of at least 80 participants to ensure statistical power). For each window, we applied the identical spectral clustering pipeline and evaluated the consistent cluster indices across each narrow age range and the full age range. All the narrow age range population yielded two distinct brain subtypes as similar as the characteristics in our main results. The assignment consistency were 90.48% (5∼8.9 yrs, N = 84), 91.89% (6∼9.9 yrs, N = 111), 98.67% (7∼10.9 yrs, N = 150), 97.71% (8∼11.9 yrs, N = 175) and 88.42% (9∼12.9 yrs, N = 190). These remarkable consistency in subgroup classification suggests that the specific impairments on brain morphology are consistently maintained across development.

The observed age difference between subgroups, though minor, could have significant implications during sensitive developmental periods. Lifespan trajectories^32^ of 34 brain regions show an early steep increase in volume, followed by a near-linear decline. The middle temporal gyrus peaks at 7.8 years, while the caudal anterior cingulate peaks at 9.2 years^32^, possibly related to the result that subgroup L is generally younger than subgroup H. However, in the CABIC cohort, we observed an inverse age relationship across both classification methods (whether using the ABIDE-classifier or independent clustering), with the subgroup L being significantly older than the H (Figure S3a and S5i). The age difference subgroups needs future studies with larger sample sizes to validate. While here, we propose some possible reasons for the ABIDE cohort to understand that these distinct structural abnormalities likely reflect differences in the timing of atypical developmental processes.

The isthmus cingulate, a key region in subgroup clustering, typically peaks in volume at 3.8 years. The lack of age-related decline in cortical volume was consistently found in subgroup H in both the ABIDE and CABIC datasets (See Supplementary Analysis and Figure S1f). However, subgroup H aligns with normative references before age 5, while subgroup L shows smaller initial volumes (Figure S1e). In subgroup H, an increase in structural covariance between the caudal middle frontal cortex (peaking at 7.75 years) and the isthmus cingulate (Figure 2c) suggests synchronized developmental trajectories. This may reflect precocious caudal middle frontal development, delayed isthmus cingulate development, or both regions following similarly abnormal patterns^22^. Correlation analyses link larger isthmus cingulate volumes in subgroup H directly with more pronounced autistic mannerisms (Figure 2a). These evidence hint at delayed development of the isthmus cingulate in subgroup H. With a normal OoS distribution (*p* = 0.06; Table 1, column 6) in this subgroup, this morphological change appears to represent a homogeneous and quantifiable pattern. In subgroup L, the parahippocampal gyrus (peaking at 10.63 years) shows reduced covariance with the entorhinal cortex (peaking at 22.67 years) but increased covariance with the lateral occipital cortex (peaking at 5.21 years). This pattern suggests a precocious development of the parahippocampal gyrus and/or the delayed lateral occipital cortex in this subgroup. Enhanced structural covariances may also reflect compensatory adaptations to mitigate structural disadvantages inherent in abnormal developmental patterns.

Recent genome-wide association studies (GWAS)^62^ reveal that cortical phenotypes exhibit distinct genetic architectures, with shared genetic variants influencing both normative brain size variation. Furthermore, significant genetic correlations between cortical expansion patterns and cognitive measures suggest multi-level mechanisms underlying neurodevelopment. These findings collectively motivate future study of ASD to build longitudinal cohorts to explore how genetically mediated cortical maturation trajectories influences different stages of development in ASD subgroups. Such approaches would bridge genotype-phenotype mapping to clinically biomarkers, advancing precision diagnostics and targeted interventions during sensitive neurodevelopmental windows.

### Reproducibility across Datasets and Implications for Biomarker Development

One key contribution of this study is the reproducibility of brain-behavior associations across the ABIDE and CABIC datasets. In our analysis, we didn’t apply on any multiple comparisons correction, three considerations may explain. First, the dual-dataset cross-validation design required consistent results across two independent cohorts (heterogeneous in culture/age), which substantially reduces false-positive risks through empirical replication. Second, as this is an exploratory target-discovery studies, overly conservative adjustments might obscure biologically meaningful results. Third, the cross-population reproducibility were prioritized in our study. We considered that uncorrected-but-replicated results had greater biological credibility than findings that were statistically corrected but failed to replicate. As the result, this cross-dataset validation confirms that the findings reflect stable neurobiological characteristics.

In subgroup H, an enlarged transverse temporal gyrus may impair precise auditory processing. This disruption can affect auditory responsiveness and the ability to integrate and understand information during social interactions. It may also damage early language perception and acquisition, particularly before a diagnosis of ASD is made. Research has found that when ASD is accompanied by cognitive learning needs, head enlargement becomes even more pronounced^45^. This suggests a shared mechanism linking brain overgrowth with impaired intellectual functioning. Alternatively, cognitive impairments might mediate the relationship between this overgrowth and ASD. Although intelligence moderates the relationship between ASD traits, transverse temporal gyrus abnormality remains a direct indicator (Figure 2b). This suggests impaired primary sensory perceptual function that is directly disrupted by abnormal sensory input. The inferior temporal gyrus (ITG) is also uniquely associated with restricted interests and repetitive behaviors, despite the low prevalence of structural abnormalities in subgroup H. This region plays a crucial role in object recognition^63^. Disruptions in these functions can lead to a tendency to focus repetitively on objects or details. While as the RRB scale in ADOS cannot depict all the dimensions of repetitive behaviors, related results need future studies focusing on more detailed investigations on RRB to validate.

We classified ASD subtypes solely based on brain morphology, assuming a causal relationship between brain structure and cognition. However, this relationship is inherently complex, thus interpreting the brain in regions goes against its interconnected nature. Although subgroup L exhibited a higher prevalence of structural abnormalities, suggesting more extensive morphological variations, we did not find reproducible brain-behavior associations between cohorts. Several factors may explain these findings. First, cognitive impairments related to structural abnormalities in subgroup L might be more sensitive to cultural differences. Second, the validity of ASD diagnoses in subgroup L might be questionable. The best predictor of a DSM-IV diagnosis has been found to be often the specific clinic attended, rather than any defining characteristic of the individual^64^. This indicates that current diagnostic tools may not capture the nuanced behaviors directly related to brain abnormalities in subgroup L. Considering global reduction in brain volume, which may result from cardiogenic etiological damage, our findings highlight the urgent need for new methods and paradigms to enable deep phenotyping and reduce the reliance on behavioral scales for differentiation. Third, heterogeneity within subgroups may also contribute to the limited reproducibility findings. Behavioral paradigms targeting brain regions with a normal distribution of OoS scores could better reveal subgroup-specific relationships between brain morphology and cognitive behavior.

In our analysis, we have rigorously controlled for site effects to minimize their impact on the results. To validate our approach, we performed independent spectral cluster analysis using only NYU site data from ABIDE. The assignment consistency compare with the main analysis reached 90.48%, demonstrating both the robustness of our clustering framework to site-specific variability and the biological validity of the identified subtypes. This high concordance further supports the clinical translatability of our neurosubtyping scheme, as it exists even when derived from diverse data sources, therefore providing a solid foundation for future multi-site biomarker development targeting subtype-specific mechanisms.

### Limitations & Future Direction

Some limitations in this study should be noted. First, previous research highlights atypical structural connectome asymmetry in ASD^65^, while we could not fully evaluate hemispheric differences within subgroups using LBCC trajectories. Second, the cross-sectional nature of the data limits causal inferences about how these abnormalities of the brain morphology arise and their relationships with ASD symptoms. Future research should thoroughly investigate brain function and cognitive differences between these subgroups while exploring how demographic, environmental, and developmental factors shape brain-based subtypes. Combining longitudinal data *invivo* and *invitro*^42^ could expand cohort diversity and provide deeper insights into the neurobiological and molecular mechanisms underlying different subtypes. When designing research cohorts to study further, focus on more representative ASD prototypes^66^ could help reduce heterogeneity within subgroups. Third, we should note that the use of ADOS rather than ADOS-2 in the CABIC cohort represents a methodological limitation that future studies with larger datasets should address.

More attention should be paid on people with more severe ASDs, such as minimally verbal autism, who remain underrepresented in brain imaging studies^67^, partly due to the challenges of collecting neuroimaging data, especially in young children. Our research strategy uses structural magnetic resonance imaging, feasible in natural sleep or sedation, with significant clinical potential. However, multimodal approaches with higher ecological validity are essential to identify neurosubtypes that include these samples. Importantly, all current findings are derived from male ASD cohorts, any clinical translations should be cautiously limited to male populations until validated in females. Therefore, future studies should expand ASD cohorts with more female representation to enable comprehensive study of sex-specific brain morphological heterogeneity.

At the same time, replication with different clustering algorithms is required to confirm the stability of these findings. For example, we employed Gaussian mixture modeling (GMM) as a validation framework to assess consistency with our main clustering results. GMM assumes the observed data are generated from a mixture of multiple distinct Gaussian distributions^68^. As the high cluster indices consistency (83.58%) across both methods, the brain morphology and cognitive behavior have the similar characteristics as spectral clustering analysis (Figure S6). Participants in subgroup L were also significantly younger than those in subgroup H (*p* = 0.03). No significant differences were observed in other comparisons between two subgroups. These replication with GMM demonstrates that our ASD neurosubtyping is not method-dependent. While more nuanced clustering approaches and strategies should be explored in the future, as disease variations may exist within the range of normal variation^69^, and multiple subpopulations may also exist in typically developing populations^70^. Finally, translational studies using animal models are essential for understanding the mechanisms underlying these different subtypes. Such studies can provide critical information on how abnormalities in brain morphology develop^71^.

### Conclusion

Linking specific brain morphology abnormalities to cognitive or behavioral impairments in ASD is challenging. Different biological mechanisms can lead to similar behavioral outcomes, complicating the identification of neural correlates. This overlap may explain the difficulty in replicating neuroimaging findings between studies or cohorts. Using the largest brain normative datasets and two large-scale cross-cultural consortiums, we applied normative modeling to disentangle the heterogeneity of brain morphology in ASD. By breaking the ASD population into smaller and more morphologically homogeneous subgroups, we identified two subtypes of morphological abnormalities. We identified significant correlations between cortical regions and autistic traits across both consortia, with particular emphasis on the isthmus cingulate cortex, as well as the transverse and inferior temporal gyri. Based on the abnormal prevalence of regional volumes, we found two structural impairment pathways that could disrupt sensory to higher cognitive functions. These findings highlight the importance of untangling mixed biological mechanisms and offer insight to develop effective subgroup-driven individualized interventions. In conclusion, our study provides a reference for elucidating the etiological mechanisms of neurodevelopmental disorders and advancing future subgroup-driven precision clinical practice.

## MATERIALS AND METHODS

### Participants and Data Preprocessing

Participants were selected from the Autism Brain Imaging Data Exchange (ABIDE) cohort^34,35^. The original dataset includes 1,833 autistic and 1,838 non-autistic individuals aged 2∼64 years from 56 sites. All diagnoses were based on the DSM-IV or DSM-V criteria. We used T1-weighted MRI data from the ABIDE (I and II) cohort. The acquisition parameters are publicly available at ABIDE website. Data quality was assessed through manual visual inspection aided by visualization outputs from (MRIQC)^72^ (version 22.0.6). We visually rated the quality of 2,451 images using a 3-class framework^73^, with “0” denoting images that suffered from gross artefacts and were considered unusable, “1” with some artefacts, but that were still considered usable, and “2” free from visible artefacts. Images with an “0” score (for example, with severe motion artifacts, segmentation errors, or structural abnormalities) were excluded. A total of 1907 (77.8%) images passed the quality control.

Due to the limited samples of female ASD participants, and the consideration that sex-specific developmental trajectories^32^ and autism-related neuroanatomical differences^74,75^ may confound data-driven population clustering, our study focuses only on male participants under 13 years old. This age range reflects our interest in brain development during childhood and early adolescence. This yielded 274 participants for downstream analyses. The participant ID list is provided in the Supplementary Materials.

Pre-processing performed through the Connectome Computation System (CCS) pipeline^76,77^. FreeSurfer (toolbox: version 6.0.0) was used for skull removal, cortical reconstruction, segmentation, and registration in the Montreal Neurological Institute (MNI) template. Each T1-weighted image was segmented into gray matter (GM), white matter (WM), and cerebrospinal fluid (CSF).

The total cortical volume (TCV) was calculated as the sum of the GM and WM volumes, following the LBCC method^32^. The mean cortical thickness (CT) was calculated as a weighted average of the thicknesses of the left and right hemispheres. The total surface area (SA) was determined by summing the surface areas of white matter of both hemispheres. Regional brain volumes for 34 bilaterally averaged cortical regions were evaluated using Desikan-Killiany parcellation^36^. To reduce site-related variability, we applied ComBat harmonization^78^. This process was implemented using (*neuroCombat*) R package.

### Normative Scoring and Clustering Analysis

To assess how the brain morphology of each participant aligned with age-standardized normative trajectories, we used the out-of-sample (OoS) centile score^32^. First, all non-ASD individuals from the site-harmonized ABIDE data set were employed to estimate cohort-specific statistical offsets. The centile scores of each brain measurement were then estimated for all individuals with ASD compared to the offset trajectory (see *Methods: Centile scoring of new MRI data* in the original LBCC work^32^). This approach takes advantage of the generalized additive models for location, scale, and shape (GAMLSS)^79^ framework to estimate statistical offsets such as mean, variance, and skewness. These offsets capture individual deviations from typical brain development. OoS scores are comparable between age and cohort. The scoring process was conducted with the publicly available code from Lifespan Charts website.

Using OoS scores from 34 brain regions as classification features, we applied spectral clustering^80^ among individuals with ASD in the ABIDE cohort. Spectral clustering was selected for its ability to detect nonlinear data structures, which are common in brain imaging datasets. Unlike *k*-means clustering, which relies on centroid-based optimization, spectral clustering transforms the data into a lower-dimensional space using the eigenvalues of a similarity matrix. Before clustering, the Jarque-Bera test was performed to determine whether OoS scores follow a normal distribution.

Clustering analysis was implemented using the *kernlab* package R. We tested clusters of 2 to 10 to identify the optimal solution. The quality of clustering was evaluated using the Silhouette Coefficient (SC)^81^, which assesses both within-cluster cohesion and between-cluster separation. SC values range from -1 to 1, with higher scores indicating better defined clusters. The optimal number of clusters was decided by the highest SC score.

### Key Brain Regions for Subgroup Classification

We developed a predictive model using machine learning with individual OoS scores from 34 regions of the brain as input characteristics. We trained a Support Vector Machine (SVM) classifier^82^ with Recursive Feature Elimination and Cross-Validation (RFECV). This was implemented using the *scikit-learn* library in Python. Combining recursive feature elimination (RFE)^83^ with cross-validation allows the model to iteratively identify the most informative features while maintaining generalizability and robustness. The data set was divided into 80% training sets and 20% testing sets. Using a linear SVM as the base estimator, RFECV removed less important features in a stepwise manner, with performance monitored through 5-fold cross-validation. This process ensured that the final feature set balanced predictive power and reduced over-fitting.

Following feature selection, we fine-tuned the SVM model by performing a grid search to optimize key hyperparameters, including regularization strength (*C*) and kernel coefficients (*γ*). The model was evaluated across different kernel functions, such as linear, polynomial, and radial basis functions, to determine the best configuration. Once optimized, the model was trained on the selected features to achieve optimal classification performance. SHapley Additive exPlanations (SHAP)^37^ were applied to quantify the contribution of each brain region to the model predictions.

### Structural Covariance Network Analysis

To investigate morphological relationships of a specific cluster, we constructed structural covariance analysis^84^ on OoS scores of each two regions. Pearson’s correlation analysis was performed on residuals obtained from linear regression models that controlled for site effects and TCV OoS scores. To enable statistical comparison, correlation coefficients were normalized using Fishers *r*-to-*z* transformation, defined as:

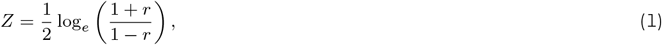

where *r* represents the Pearson correlation coefficient between two regions. The difference (cluster i/ii vs. normal control) in transformed correlation coefficients was calculated using:

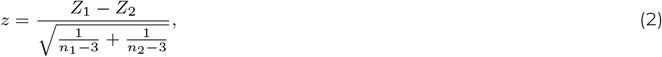

where *n*_1_ and *n*_2_ are the two sample sizes being compared. The resulting *z*-values were converted to *p*-values using the cumulative distribution function of the standard normal distribution.

### Cluster Differences and Brain-Behavior Analysis

We investigated differences between clusters and demographic, technical and clinical variables. For categorical variables, such as subtypes of perinatal developmental disorders (PDD), scanner models, and manufacturers, Chi-square tests were employed. For data collection sites, Fishers exact test was used with Monte Carlo simulations to approximate *p* values. To ensure robust analysis, sites with fewer than 10 participants were excluded. For other continuous variables, statistical methods were selected based on data distribution: the Wilcoxon rank sum test was used for nonnormally distributed data, and the two-sample *t*-test for data shows normal distribution. The effect sizes were calculated using Cramér’s *V* for categorical variables and Cohens *d* for continuous variables. Statistical significance was set at *p <* 0.05.

To further explore brain-behavior relationships within clusters, we performed correlation analyses using Full-Scale Intelligence Quotient (FIQ), ADOS-2 (Social Affect and Restricted and Repetitive Behaviors, RRB), ADI-R (Social Interaction and RRB), and SRS subscales. Pearson’s correlation was applied to most measures, while Spearman’s correlation was used for ADOS-2 RRB scores due to their limited range. In these analyzes, the effects of the site and the OoS scores of TCV were controlled. Statistical significance was set at *p <* 0.05, and no correction for multiple comparisons was applied.

For each group, we also examined partial correlations to investigate structural covariance between 34 brain regions and their association with cognitive behaviors. For each participant, the partial covariance scores of two regions were calculated as

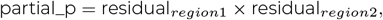

The resulting partial covariance scores were then correlated with the same measurements with the same correlated methods. Finally, we performed mediation analyzes to examine whether FIQ mediates the relationships between brain morphology and cognitive behaviors within clusters. Linear models were used to estimate direct and indirect effects, with adjustments for site and OoS scores of TCV. All brain morphology and cognitive behavior variables were standardized to ensure comparability among participants. The significance of indirect effects was assessed using the Sobel test. As the brain abnormalities are detectable in the early development^40^ before the emergence of autistic behaviors^85^, we built a mediation model to examine the brain’s effect on cognitive behaviors. To provide a more comprehensive account, we also analyzed the reverse mediation model.

### Detection of Reproducible Results Between Datasets

We applied the ABIDE-based classifier to an independent dataset from the China Autism Brain Imaging Consortium (CABIC)^33^, which is a multicenter collaboration among clinical and research institutions in China. It is designed to collect neuroimaging, demographic, and behavioral data on children and adolescents with ASD. Diagnoses in the CABIC cohort were based on DSM-IV, DSMV, or ICD-11 criteria and validated using ADOS and ADI-R. The dataset includes 2,656 participants, including 1,503 ASD individuals and 1,153 typically developing children. Detailed MRI acquisition parameters are publicly accessible at CABIC Site. There were in total 968 ASD males (4.56±1.69) and 526 (7.37±2.73) typically controls included into analysis.

CABIC participants, aged 1 to 13 years, tend to exhibit more severe ASD symptoms, resulting in lower variability within clusters. Although this homogeneity is clinically valuable, it poses challenges in detecting subtle brain-behavior relationships. In contrast, the ABIDE cohort provides greater behavioral differences, enhancing the ability to identify nuanced patterns in brain structure. By leveraging these complementary datasets, our goal was to identify results that are reproducible across both cohorts. We repeated the statistical, structural covariance, and correlation analyses on CABIC cohort. However, the limited FIQ data in CABIC restricted the replication of mediation analysis with sufficient statistical power. Therefore, mediation analysis was conducted on the ABIDE cohort only.

In this study, we applied a SVM classifier trained on the ABIDE cohort on the CABIC cohort. To evaluate the consistency between this transfer approach and independent clustering approach, we replicated the identical analytical process directly on ASD participants from CABIC cohort (See details in the Supplementary Information). Optimal cluster solutions was also two (Figure S5a). Among 968 participants, 929 (95.97%) exhibited consistent cluster indices across both methods. This high concordance highlights the robustness of our spectral clustering framework across heterogeneous datasets.

## ACKNOWLEDGMENTS

The STI 2030 - the major projects of the Brain Science and Brain-Inspired Intelligence Technology (2021ZD0200500), the Start-up Funds for Leading Talents at Beijing Normal University, the National Natural Science Foundation of China (82102134, 82322035, 62273076), the Key-Area Research and Development Program of Guangdong Province (2019B030335001), the Beijing Municipal Science and Technology Commission (Z161100002616023, Z181100001518003), the Major Project of National Social Science Foundation of China (20&ZD296), the CAS-NWO Programme (153111KYSB20160020), the National Basic Research (973) Program (2015CB351702), the Chinese Academy of Sciences Key Research Program (KSZD-EW-TZ-002) and the Major Fund for International Collaboration of National Natural Science Foundation of China (81220108014) provide funding supports.

## AUTHOR CONTRIBUTIONS

Conception and Design: X-R.F., X-N.Z. Data Analysis: X-R.F. MRI Data Preprocessing: Y.H., CABIC. Initial Drafting of the Manuscript: X-R.F. Critical Review and Editing of the Manuscript: All authors contributed to the critical review and editing of the manuscript.

## DATA AVAILABILITY

The datasets analyzed in this study are publicly available. The ABIDE dataset can be accessed from the Autism Brain Imaging Data Exchange repository at http://fcon_1000.projects.nitrc.org/indi/abide. The CABIC dataset is available upon request through the CABIC Data Sharing Initiative at https://php.bdnilab.com. The key data we used in this study are the OoS centile scores we calculated for each participant with ABIDE and CABIC. To foster researcher on replication and extended exploration, we shared these data through the *Chinese Color Nest Data Community: Fostering Lifespan Development of Brain-Mind Health*. Data have been deposited into CSV files^86^ at https://doi.org/10.57760/sciencedb.24587, and they are accessible upon the requests according to the instructions described on this page.

## CODE AVAILABILITY

All code used in the analyses is available at https://github.com/XueruFan/ASD-Project. For detailed information on how to use these codes and replicate this study step-by-step, please see the explanation in the README.md file.

## For the Lifespan Brain Chart Consortium (LBCC)

Chris Adamson^4,5^, Sophie Adler^6^, Aaron F. Alexander-Bloch^7,8,9^, Evdokia Anagnostou^10,11^, Kevin M. Anderson^12^, Ariosky Areces-Gonzalez^13,14^, Duncan E. Astle^15^, Bonnie Auyeung^16,17^, Muhammad Ayub^18,19^, Jong Bin Bae^20^, Gareth Ball^4,21^, Simon Baron-Cohen^17,22^, Richard Beare^4,5^, Saashi A. Bedford^17^, Vivek Benegal^23^, Richard A.I. Bethlehem^17,24^, Frauke Beyer^25^, John Blangero^26^, Manuel Blesa Cábez^27^, James P. Boardman^27^, Matthew Borzage^28^, Jorge F. Bosch-Bayard^29,30^, Niall Bourke^31^, Edward T. Bullmore^32^, Vince D. Calhoun^33^, Mallar M. Chakravarty^34,30^, Christina Chen^35^, Casey Chertavian^9^, Gaël Chetelat^36^, Yap S. Chong^37,38^, Aiden Corvin^39^, Manuela Costantino^40,41^, Eric Courchesne^42,43^, Fabrice Crivello^44^, Vanessa L. Cropley^45^, Jennifer Crosbie^46^, Nicolas Crossley^47,48^, Marion Delarue^36^, Richard Delorme^49,50^, Sylvane Desrivieres^51^, Gabriel Devenyi^52,53^, Maria A. Di Biase^45,54^, Ray Dolan^55,56,57^, Kirsten A. Donald^58,59^, Gary Donohoe^60^, Lena Dorfschmidt^7,8,9^, Katharine Dunlop^61^, Anthony D Edwards^62,63,64^, Jed T. Elison^65^, Cameron T. Ellis^12,66^, Jeremy A. Elman^67^, Lisa Eyler^68,69^, Damien A. Fair^65^, Paul C. Fletcher^70,71^, Peter Fonagy^72,73^, Carol E. Franz^74^, Lidice Galan-Garcia^75^, Ali Gholipour^76^, Jay Giedd^77,78^, John H. Gilmore^79^, David C. Glahn^80,81^, Ian M. Goodyer^32^, P. E. Grant^82^, Nynke A. Groenewold^83,59^, Shreya Gudapati^7,8,9^, Faith M. Gunning^84^, Raquel E. Gur^7,9^, Ruben C. Gur^7,85^, Christopher F. Hammill^46,86^, Oskar Hansson^87,88^, Trey Hedden^89,90^, Andreas Heinz^91^, Richard N. Henson^15,92^, Katja Heuer^93^, Jacqueline Hoare^94^, Bharath Holla^95,96^, Avram J. Holmes^97^, Hao Huang^98^, Jonathan Ipser^99^, Clifford R. Jack Jr^100^, Andrea P. Jackowski^101,102^, Tianye Jia^103,104,105^, David T. Jones^106,107^, Peter B. Jones^32,108^, Rene S Kahn^109,110^, Hasse Karlsson^111,112^, Linnea Karlsson^111,112^, Ryuta Kawashima^113^, Elizabeth A. Kelley^114^, Silke Kern^115,116^, Ki-Woong Kim^117,118,119,120^, Manfred G. Kitzbichler^32^, William S. Kremen^67^, François Lalonde^121^, Brigitte Landeau^36^, Jason Lerch^122,123,124^, John D. Lewis^125^, Jiao Li^126^, Wei Liao^126^, Conor Liston^127^, Michael V. Lombardo^128,17^, Jinglei Lv^129,45^, Travis T. Mallard^130^, Machteld Marcelis^131^, Samuel R. Mathias^80^, Bernard Mazoyer^44,132^, Philip McGuire^133^, Michael J. Meaney^134^, Andrea Mechelli^133^, Bratislav Misic^135^, Sarah E Morgan^136,137,138^, David Mothersill^139,140,141^, Cynthia Ortinau^142^, Rik Ossenkoppele^143,144^, Minhui Ouyang^98^, Lena Palaniyappan^145^, Leo Paly^36^, Pedro M. Pan^146,147^, Christos Pantelis^148,149,150^, Min Tae M. Park^151^, Tomas Paus^152,153^, Zdenka Pausova^46,154^, Deirel Paz-Linares^13,155^, Alexa Pichet Binette^156,157^, Karen Pierce^158^, Xing Qian^159^, Anqi Qiu^160^, Armin Raznahan^121^, Timothy Rittman^161^, Amanda Rodrigue^80^, Caitlin K. Rollins^162,163^, Rafael Romero-Garcia^32,164^, Lisa Ronan^32^, Monica D. Rosenberg^165^, David H. Rowitch^166^, Giovanni A. Salum^167,168^, Theodore D. Satterthwaite^169,7^, H. Lina Schaare^170,171^, Jenna Schabdach^7,8,9^, Russell J. Schachar^46^, Michael Schöll^172,173,174^, Aaron P. Schultz^175,176,81^, Jakob Seidlitz^7,8,9^, David Sharp^177,178^, Russell T. Shinohara^35,179^, Ingmar Skoog^115,116^, Christopher D. Smyser^180^, Reisa A. Sperling^175,181,81^, Dan J. Stein^182^, Aleks Stolicyn^183^, John Suckling^32,184^, Gemma Sullivan^27^, Benjamin Thyreau^113^, Roberto Toro^93,185^, Nicolas Traut^186,187^, Kamen A. Tsvetanov^161,188^, Nicholas B. Turk-Browne^12,189^, Jetro J. Tuulari^190,191,192^, Christophe Tzourio^193^, Étienne Vachon-Presseau^194,195,196^, Mitchell J. Valdes-Sosa^75^, Pedro A. Valdes-Sosa^197,198^, Sofie L. Valk^199^, Therese van Amelsvoort^200^, Simon N. Vandekar^201,202^, Lana Vasung^203^, Petra E Vértes^32,204^, Lindsay W. Victoria^84^, Sylvia Villeneuve^157,205,156^, Arno Villringer^25,206^, Jacob W. Vogel^169,7^, Konrad Wagstyl^207^, Yin-Shan Wang^1,208,209,210^, Simon K. Warfield^76^, Varun Warrier^211^, Eric Westman^212^, Margaret L. Westwater^32^, Heather C. Whalley^183,213^, Simon R. White^32,214^, A. Veronica Witte^206,25,215^, Ning Yang^1,208,209,210^, B.T. Thomas Yeo^216,217,218,219^, Hyuk Jin Yun^220^, Andrew Zalesky^221^, Heather J Zar^83,222^, Anna Zettergren^115^, Juan H. Zhou^159,223,216^, Hisham Ziauddeen^32,224,^ Dabriel Zimmerman^7,8,9^, Andre Zugman^226,227,147^, Xi-Nian Zuo^1,208,209,228^

### LBCC Affiliations

^4^Developmental Imaging, Murdoch Children’s Research Institute, Melbourne, Victoria, Australia

^5^Department of Medicine, Monash University, Melbourne, Victoria, Australia

^6^UCL Great Ormond Street Institute for Child Health, 30 Guilford St, Holborn, London WC1N 1EH

^7^Department of Psychiatry, University of Pennsylvania, Philadelphia, PA 19104

^8^Department of Child and Adolescent Psychiatry and Behavioral Science, The Children’s Hospital of Philadelphia, Philadelphia, PA 19104

^9^Lifespan Brain Institute, The Children’s Hospital of Philadelphia, Philadelphia, PA 19104

^10^Department of Pediatrics University of Toronto

^11^Holland Bloorview Kids Rehabilitation Hospital, Toronto, Canada

^12^Department of Psychology, Yale University, New Haven, CT, USA

^13^The Clinical Hospital of Chengdu Brain Science Institute, MOE Key Lab for NeuroInformation, University of Electronic Science and Technology of China, No. 2006, Xiyuan Ave., West Hi-Tech Zone, Chengdu, 611731, China

^14^University of Pinar del Río Hermanos Saiz Montes de Oca, Cuba

^15^MRC Cognition and Brain Sciences Unit, University of Cambridge, Cambridge UK

^16^Department of Psychology, School of Philosophy, Psychology and Language Sciences, University of Edinburgh, Edinburgh, United Kingdom

^17^Autism Research Centre, Department of Psychiatry, University of Cambridge, Cambridge, CB2 0SZ, UK

^18^Queen’s University, Department of Psychiatry, Centre for Neuroscience Studies, Kingston, Ontario, Canada

^19^University College London, Mental Health Neuroscience Research Department, Division of Psychiatry, London UK

^20^Department of Neuropsychiatry, Seoul National University Bundang Hospital, Seongnam, Korea

^21^Department of Paediatrics, University of Melbourne, Melbourne, Victoria, Australia

^22^Cambridge Lifetime Asperger Syndrome Service (CLASS), Cambridgeshire and Peterborough NHS Foundation Trust, Cambridge, United Kingdom

^23^Centre for Addiction Medicine, National Institute of Mental Health and Neurosciences (NIMHANS), Bengaluru, India 560029

^24^Brain Mapping Unit, Department of Psychiatry, University of Cambridge, Cambridge, CB2 0SZ, UK

^25^Department of Neurology, Max Planck Institute for Human Cognitive and Brain Sciences, Leipzig, 04103, Germany

^26^Department of Human Genetics, South Texas Diabetes and Obesity Institute, University of Texas Rio Grande Valley

^27^MRC Centre for Reproductive Health, University of Edinburgh, UK

^28^Fetal and Neonatal Institute, Division of Neonatology, Children’s Hospital Los Angeles, Department of Pediatrics, Keck School of Medicine, University of Southern California, Los Angeles, California USA

^29^McGill Centre for Integrative Neuroscience, Ludmer Centre for Neuroinformatics and Mental Health, Montreal Neurological Institute

^30^McGill University

^31^Department of Brain Sciences, Imperial College London, London UK & Care Research & Technology Centre, UK Dementia Research Institute

^32^Department of Psychiatry, University of Cambridge, Cambridge, CB2 0SZ, UK

^33^Tri-institutional Center for Translational Research in Neuroimaging and Data Science, Georgia State University, Georgia Institute of Technology, and Emory University, Atlanta, GA, USA

^34^Computational Brain Anatomy (CoBrA) Laboratory, Cerebral Imaging Centre, Douglas Mental Health University Institute

^35^Penn Statistics in Imaging and Visualization Center, Department of Biostatistics, Epidemiology, and Informatics, Perelman School of Medicine, University of Pennsylvania, Philadelphia, PA, USA

^36^Normandie Univ, UNICAEN, INSERM, U1237, PhIND “Physiopathology and Imaging of Neurological Disorders”, Institut Blood and Brain @ Caen-Normandie, Cyceron, 14000 Caen, France

^37^Singapore Institute for Clinical Sciences, Agency for Science, Technology and Research, Singapore

^38^Department of Obstetrics and Gynaecology, Yong Loo Lin School of Medicine, National University of Singapore, Singapore

^39^Department of Psychiatry, Trinity College, Dublin, Ireland

^40^Cerebral Imaging Centre, Douglas Mental Health University Institute, Verdun, Canada

^41^Undergraduate program in Neuroscience, McGill University, Montreal, Canada

^42^Department of Neuroscience, University of California, San Diego, San Diego, CA 92093, USA

^43^Autism Center of Excellence, University of California, San Diego, San Diego, CA 92037, USA

^44^Institute of Neurodegenerative Disorders, CNRS UMR5293, CEA, University of Bordeaux

^45^Melbourne Neuropsychiatry Centre, University of Melbourne, Melbourne, Australia

^46^The Hospital for Sick Children, Toronto, Canada

^47^Department of Psychiatry, School of Medicine, Pontificia Universidad Católica de Chile, Diagonal Paraguay 362, Santiago 8330077, Chile

^48^Department of Psychiatry, University of Oxford OX3 7JX

^49^Child and Adolescent Psychiatry Department, Robert Debré University Hospital, AP-HP, F-75019, Paris France

^50^Human Genetics and Cognitive Functions, Institut Pasteur, F-75015, Paris France

^51^Social, Genetic and Developmental Psychiatry Centre, Institute of Psychiatry, Psychology & Neuroscience, King’s College London, London, United Kingdom

^52^Cerebral Imaging Centre, Douglas Mental Health University Institute, Montreal, QC, Canada, McGill Department of Psychiatry, Montreal, QC, Canada

^53^Department of Psychiatry, McGill University, Montreal, QC, Canada

^54^Department of Psychiatry, Brigham and Womens Hospital, Harvard Medical School, Boston, Massachusetts, United States

^55^Max Planck UCL Centre for Computational Psychiatry and Ageing Research, University College London, London, UK

^56^Wellcome Centre for Human Neuroimaging, University College London, London, UK

^57^Wellcome Centre for Human Neuroimaging, 12 Queen Square, London WC1N 3AR

^58^Division of Developmental Paediatrics, Department of Paediatrics and Child Health, Red Cross War Memorial Children’s Hospital, Klipfontein Road/Private Bag, Rondebosch, 7700/7701, Cape Town, South Africa

^59^Neuroscience Institute, University of Cape Town, Cape Town, South Africa

^60^Center for Neuroimaging, Cognition & Genomics (NICOG), School of Psychology, National University of Ireland Galway, Galway, Ireland

^61^Weil Family Brain and Mind Research Institute, Department of Psychiatry, Weill Cornell Medicine

^62^Centre for the Developing Brain, King’s College London, London, UK

^63^Evelina London Children’s Hospital

^64^MRC Centre for Neurodevelopmental Disorders, London

^65^Institute of Child Development, Department of Pediatrics, Masonic Institute for the Developing Brain, University of Minnesota, Minneapolis, MN, United States

^66^Haskins Laboratories, New Haven, CT, USA

^67^Department of Psychiatry, Center for Behavior Genetics of Aging, University of California, San Diego, La Jolla, CA

^68^Desert-Pacific Mental Illness Research Education and Clinical Center, VA San Diego Healthcare, San Diego, CA, USA

^69^Department of Psychiatry, University of California San Diego, Los Angeles, CA, USA

^70^Department of Psychiatry, University of Cambridge, and Wellcome Trust MRC Institute of Metabolic Science, Cambridge Biomedical Campus, Cambridge, United Kingdom

^71^Cambridgeshire and Peterborough NHS Foundation Trust

^72^Department of Clinical, Educational and Health Psychology, University College London, London, UK

^73^Anna Freud National Centre for Children and Families, London UK

^74^Department of Psychiatry, Center for Behavior Genetics of Aging, University of California, San Diego, La Jolla, CA 92093

^75^Cuban Center for Neuroscience, La Habana, Cuba

^76^Computational Radiology Laboratory, Boston Childrens Hospital, Boston, MA 02115

^77^Department of Child and Adolescent Psychiatry, University of California, San Diego, San Diego, CA 92093, USA

^78^Department of Psychiatry, University of California San Diego, San Diego, CA, USA

^79^Department of Psychiatry, University of North Carolina, Chapel Hill, NC, USA

^80^Department of Psychiatry, Boston Childrens Hospital and Harvard Medical School, Boston, MA 02115

^81^Harvard Medical School, Boston, MA 02115

^82^Division of Newborn Medicine and Neuroradiology, Fetal Neonatal Neuroimaging and Developmental Science Center, Boston Childrens Hospital, Harvard Medical School, Boston, MA 02115, USA

^83^Department of Paediatrics and Child Health, Red Cross War Memorial Children’s Hospital, SA-MRC Unit on Child & Adolescent Health, University of Cape Town, South Africa

^84^Weill Cornell Institute of Geriatric Psychiatry, Department of Psychiatry, Weill Cornell Medicine

^85^Lifespan Brain Institute, The Children’s Hospital of Philadelphia, Philadelphia, PA 19105

^86^Mouse Imaging Centre, Toronto, Canada

^87^Clinical Memory Research Unit, Department of Clinical Sciences Malmö, Lund University, Malmö, Sweden

^88^Memory Clinic, Skåne University Hospital, Malmö, Sweden

^89^Department of Neurology, Icahn School of Medicine at Mount Sinai, New York, NY 10029, USA

^90^Athinoula A. Martinos Center for Biomedical Imaging, Department of Radiology, Massachusetts General Hospital, Harvard Medical School, Boston, MA 02129, USA

^91^Department of Psychiatry and Psychotherapy, Charite University Hospital Berlin, Berlin, Germany

^92^Department of Psychiatry, University of Cambridge, Cambridge, UK

^93^Institut Pasteur, Université Paris Cité, Unité de Neuroanatomie Appliquée et Théorique, F-75015 Paris, France

^94^Department of Psychiatry, University of Cape Town, Cape Town, South Africa

^95^Department of Integrative Medicine, NIMHANS, Bengaluru-560029, India

^96^Accelerator Program for Discovery in Brain disorders using Stem cells (ADBS), Department of Psychiatry, NIMHANS, Bengaluru-560029, India

^97^Department of Psychiatry, Brain Health Institute, Rutgers University, Piscataway, NJ, USA

^98^Department of Radiology, Children’s Hospital of Philadelphia and University of Pennsylvania, Philadelphia, PA 19104

^99^Department of Psychiatry and Mental Health, Clinical Neuroscience Institute, University of Cape Town

^100^Department of Radiology, Mayo Clinic, Rochester, MN 55905, USA

^101^Department of Psychiatry, Universidade Federal de São Paulo

^102^National Institute of Developmental Psychiatry, CNPq

^103^Institute of Science and Technology for Brain-Inspired Intelligence, Fudan University, Shanghai, 200433, China

^104^Key Laboratory of Computational Neuroscience and BrainInspired Intelligence (Fudan University), Ministry of Education, Shanghai, China

^105^Centre for Population Neuroscience and Precision Medicine (PONS), Institute of Psychiatry, Psychology and Neuroscience, SGDP Centre, King’s College London, London SE5 8AF, UK

^106^Department of Neurology, Mayo Clinic, Rochester, MN, USA

^107^Department of Radiology, Mayo Clinic, Rochester, MN, USA

^108^Cambridgeshire and Peterborough NHS Foundation Trust, Huntingdon, United Kingdom

^109^Department of Psychiatry, Icahn School of Medicine at Mount Sinai, New York, NY, USA

^110^Department of Psychiatry, Icahn School of Medicine, Mount Sinai, New York, USA

^111^Department of Clinical Medicine, Department of Psychiatry and Turku Brain and Mind Center, FinnBrain Birth Cohort Study, University of Turku and Turku University Hospital, Turku, Finland

^112^Centre for Population Health Research, Turku University Hospital and University of Turku, Turku, Finland

^113^Institute of Development, Aging and Cancer, Tohoku University, Seiryocho, Aobaku, Sendai 980-8575, Japan

^114^Queen’s University, Departments of Psychology and Psychiatry, Centre for Neuroscience Studies, Kingston, Ontario, Canada

^115^Neuropsychiatric Epidemiology Unit, Department of Psychiatry and Neurochemistry, Institute of Neuroscience and Physiology, the Sahlgrenska Academy, Centre for Ageing and Health (AGECAP) at the University of Gothenburg, Sweden

^116^Region Västra Götaland, Sahlgrenska University Hospital, Psychiatry, Cognition and Old Age Psychiatry Clinic, Gothenburg, Sweden

^117^Department of Brain and Cognitive Sciences, Seoul National University College of Natural Sciences, Seoul, Republic of Korea

^118^Department of Neuropsychiatry, Seoul National University Bundang Hospital, Seongnam, Republic of Korea ^119^Department of Psychiatry, Seoul National University College of Medicine, Seoul, Republic of Korea

^120^Department of Brain and Cognitive Science, Seoul National University College of Natural Sciences

^121^Section on Developmental Neurogenomics, Human Genetics Branch, National Institute of Mental Health, Bethesda, MD, USA

^122^Department of Medical Biophysics, University of Toronto, Toronto, ON, Canada

^123^Mouse Imaging Centre, The Hospital for Sick Children, Toronto, ON, Canada

^124^Wellcome Centre for Integrative Neuroimaging, FMRIB, Nuffield Department of Clinical Neuroscience, University of Oxford, Oxford, UK

^125^Montreal Neurological Institute, McGill University, Montreal, Canada

^126^The Clinical Hospital of Chengdu Brain Science Institute, University of Electronic Science and Technology of China, Chengdu 611731, China

^127^Department of Psychiatry and Brain and Mind Research Institute, Weill Cornell Medicine

^128^Laboratory for Autism and Neurodevelopmental Disorders, Center for Neuroscience and Cognitive Systems @UniTn, Istituto Italiano di Tecnologia, Rovereto, Italy

^129^School of Biomedical Engineering & Brain and Mind Centre, The University of Sydney, Sydney, NSW, Australia

^130^Department of Psychology, University of Texas, Austin, Texas 78712, USA

^131^Department of Psychiatry and Neuropsychology, School of Mental Health and Neuroscience, EURON, Maastricht University Medical Centre, PO Box 616, 6200 MD, Maastricht, the Netherlands; Institute for Mental Health Care Eindhoven (GGzE), Eindhoven, the Netherlands

^132^Bordeaux University Hospital

^133^Professor, Department of Psychosis Studies, Institute of Psychiatry, Psychology and Neuroscience, King’s College London, UK

^134^Ludmer Centre for Neuroinformatics and Mental Health, Douglas Mental Health University Institute, McGill University, Montreal, Quebec, Canada; Singapore Institute for Clinical Sciences, Singapore

^135^McConnell Brain Imaging Centre, Montreal Neurological Institute, McGill University, Montreal, QC H3A 2B4, Canada

^136^Department of Computer Science and Technology, University of Cambridge, Cambridge CB3 0FD, United Kingdom

^137^Department of Psychiatry, University of Cambridge, Cambridge CB2 0SZ, United Kingdom

^138^The Alan Turing Institute, London, NW1 2DB

^139^Department of Psychology, School of Business, National College of Ireland, Dublin, Ireland

^140^School of Psychology & Center for Neuroimaging and Cognitive Genomics, National University of Ireland Galway, Galway, Ireland

^141^Department of Psychiatry, Trinity College Dublin, Dublin, Ireland

^142^Department of Pediatrics, Washington University in St. Louis, St. Louis, Missouri, United States

^143^Alzheimer Center Amsterdam, Department of Neurology, Amsterdam Neuroscience, Vrije Universiteit Amsterdam, Amsterdam UMC, Amsterdam, The Netherlands

^144^Lund University, Clinical Memory Research Unit, Lund, Sweden

^145^Robarts Research Institute & The Brain and Mind Institute, University of Western Ontario,London,Ontario,Canada

^146^Department of Psychiatry, Federal University of Sao Poalo (UNIFESP)

^147^National Institute of Developmental Psychiatry for Children and Adolescents (INPD), Brazil

^148^Melbourne Neuropsychiatry Centre, Department of Psychiatry, The University of Melbourne and Melbourne Health, Carlton South, Victoria, Australia

^149^Melbourne School of Engineering, The University of Melbourne, Parkville, Victoria, Australia

^150^Florey Institute of Neuroscience and Mental Health, Parkville, VIC, Australia

^151^Department of Psychiatry, Schulich School of Medicine and Dentistry, Western University, London, ON, Canada

^152^Department of Psychiatry, Faculty of Medicine and Centre Hospitalier Universitaire Sainte-Justine, University of Montreal, Montreal, Quebec, Canada

^153^Departments of Psychiatry and Psychology, University of Toronto, Toronto, ON, Canada

^154^Departments of Physiology and Nutritional Sciences, University of Toronto, Toronto, Canada

^155^Cuban Neuroscience Center, Havana, Cuba

^156^Department of Psychiatry, Faculty of Medicine, McGill University, Montreal, Qc, H3A 1Y2, Canada

^157^Douglas Mental Health University Institute, Montreal, Qc, H4H 1R3, Canada

^158^Department of Neurosciences, University of California, San Diego La Jolla, CA, USA

^159^Center for Sleep and Cognition, Yong Loo Lin School of Medicine, National University of Singapore, Singapore

^160^Department of Biomedical Engineering, The N.1 Institute for Health, National University of Singapore ^161^Department of Clinical Neurosciences, University of Cambridge, Cambridge UK

^162^Department of Neurology, Harvard Medical School

^163^Department of Neurology, Boston Childrens Hospital, Boston, MA 02115

^164^Instituto de Biomedicina de Sevilla (IBiS) HUVR/CSIC/Universidad de Sevilla, Dpto. de Fisiología Médica y Biofísica, Spain

^165^Department of Psychology, Neuroscience Institute, University of Chicago

^166^Department of Paediatrics and Wellcome-MRC Cambridge Stem Cell Institute, University of Cambridge, Hills Road, Cambridge, UK

^167^Department of Psychiatry, Universidade Federal do Rio Grande do Sul (UFRGS)

^168^National Institute of Developmental Psychiatry (INPD)

^169^Lifespan Informatics & Neuroimaging Center, University of Pennsylvania, Philadelphia, PA 19104

^170^Otto Hahn Group Cognitive Neurogenetics, Max Planck Institute for Human Cognitive and Brain Sciences, Leipzig, Germany

^171^Institute of Neuroscience and Medicine (INM-7: Brain and Behaviour), Research Centre Juelich, Juelich, Germany

^172^Wallenberg Centre for Molecular and Translational Medicine, University of Gothenburg, Gothenburg, Sweden

^173^Department of Psychiatry and Neurochemistry, University of Gothenburg, Sweden

^174^Dementia Research Centre, Queen’s Square Institute of Neurology, University College London, UK

^175^Harvard Aging Brain Study, Department of Neurology, Massachusetts General Hospital, Boston, MA 02114

^176^Athinoula A. Martinos Center for Biomedical Imaging, Department of Radiology, Massachusetts General Hospital, Charlestown, MA 02129, USA

^177^Department of Brain Sciences, Imperial College London, London UK

^178^Care Research & Technology Centre, UK Dementia Research Institute

^179^Center for Biomedical Image Computing and Analytics, Department of Radiology, Perelman School of Medicine, University of Pennsylvania, Philadelphia, PA, USA

^180^Departments of Neurology, Pediatrics, and Radiology, Washington University School of Medicine, St. Louis, United States

^181^Center for Alzheimer Research and Treatment, Department of Neurology, Brigham and Womens Hospital, Boston, MA 02115

^182^SA MRC Unit on Risk & Resilience in Mental Disorders, Dept of Psychiatry and Neuroscience Institute, University of Cape Town, Cape Town, South Africa

^183^Division of Psychiatry, Centre for Clinical Brain Sciences, University of Edinburgh, UK

^184^Cambridge and Peterborough Foundation NHS Trust

^185^Université de Paris, Paris, France

^186^Department of Neuroscience, Institut Pasteur, Paris, France

^187^Center for Research and Interdisciplinarity (CRI), Université Paris Descartes, Paris, France

^188^Department of Psychology, University of Cambridge, Cambridge, UK

^189^Wu Tsai Institute, Yale University, New Haven, CT, USA

^190^Department of Clinical Medicine, Department of Psychiatry, FinnBrain Birth Cohort Study, University of Turku, Turku, Finland

^191^Department of Clinical Medicine, University of Turku, Turku Finland

^192^Turku Collegium for Science, Medicine and Technology, University of Turku, Turku, Finland

^193^Univ. Bordeaux, Inserm, Bordeaux Population Health Research Center, U1219, CHU Bordeaux, F-33000 Bordeaux, France

^194^Faculty of Dental Medicine and Oral Health Sciences, McGill University, Montreal, Qc, H3A 1G1, Canada

^195^Faculty of Dentistry, McGill University, Montreal, Qc, H3A 1G1, Canada

^196^Alan Edwards Centre for Research on Pain (AECRP), McGill University, Montreal, Qc, H3A 1G1, Canada

^197^Joint China-Cuba Lab,University of Electronic Science and Technology, Chengdu China/Cuban Center for Neuroscience, La Habana, Cuba

^198^University of Electronic Science and Technology of China/Cuban Center for Neuroscience

^199^Institute for Neuroscience and Medicine 7, Forschungszentrum Juelich; Max Planck Institute for Human Cognitive and Brain Sciences

^200^Department of Psychiatry & Neurosychology, Maastricht University, Maastricht, The Netherlands

^201^Department of Biostatistics, Vanderbilt University, Nashville, Tennessee, USA

^202^Department of Biostatistics, Vanderbilt University Medical Center, Nashville, Tennessee, USA

^203^Division of Newborn Medicine, Fetal Neonatal Neuroimaging and Developmental Science Center, Department of Pediatrics, Boston Childrens Hospital, Boston, MA 02115

^204^The Alan Turing Institute, London NW1 2DB, UK

^205^McConnell Brain Imaging Center, Montreal Neurological Institute, McGill University, Montreal, Quebec, Canada

^206^Clinic for Cognitive Neurology, University of Leipzig Medical Center, Leipzig, 04103, Germany

^207^Wellcome Centre for Human Neuroimaging, Institute of Neurology, University College London, WC1N 3AR

^208^Developmental Population Neuroscience Research Center, IDG/McGovern Institute for Brain Research, Beijing Normal University, Beijing 100875, China

^209^National Basic Science Data Center, Beijing 100190, China

^210^Research Center for Lifespan Development of Brain and Mind, Institute of Psychology, Chinese Academy of Sciences, Beijing 100101, China

^211^Department of Psychiatry, University of Cambridge, Cambridge, CB2 0SZ, UK

^212^Division of Clinical Geriatrics, Center for Alzheimer Research, Department of Neurobiology, Care Sciences and Society, Karolinska Institutet, Stockholm, Sweden

^213^Generation Scotland, University of Edinburgh

^214^MRC Biostatistics Unit, University of Cambridge, Cambridge, England

^215^Faculty of Medicine, CRC 1052 Obesity Mechanisms, University of Leipzig, Leipzig, 04103, Germany

^216^Department of Electrical and Computer Engineering, National University of Singapore, Singapore

^217^Centre for Sleep & Cognition and Centre for Translational MR Research, Yong Loo Lin School of Medicine, National University of Singapore, Singapore

^218^N.1 Institute for Health & Institute for Digital Medicine, National University of Singapore, Singapore

^219^Integrative Sciences and Engineering Programme (ISEP), National University of Singapore, Singapore

^220^Fetal Neonatal Neuroimaging and Developmental Science Center, Division of Newborn Medicine, Boston Childrens Hospital, Harvard Medical School, Boston, MA 02115, USA

^221^Melbourne Neuropsychiatry Centre, University of Melbourne, Melbourne, Australia; Department of Biomedical Engineering, University of Melbourne, Melbourne, Australia

^222^SAMRC Unit on Child & Adolescent Health, University of Cape Town, South Africa

^223^Center for Translational Magnetic Resonance Research, Yong Loo Lin School of Medicine, National University of Singapore, Singapore

^224^Wellcome Trust-MRC Institute of Metabolic Science, University of Cambridge, Cambridge, CB2 0SZ

^225^Cambridgeshire and Peterborough Foundation Trust, Cambridge, CB21 5EF

^226^National Institute of Mental Health (NIMH), National Institutes of Health (NIH), Bethesda, Maryland, USA

^227^Department of Psychiatry, Escola Paulista de Medicina, São Paulo, Brazil

^228^Research Center for Lifespan Development of Brain and Mind, Institute of Psychology, Chinese Academy of Sciences, Beijing 100101, China; Key Laboratory of Brain and Education, School of Education Science, Nanning Normal University, Nanning 530001, China

## For the China Autism Brain Imaging Consortium (CABIC)

Miaoshui Bai^229^, Jinhua Cai^230^, Kelong Cai^231,232^, Doudou Cao^233^, Xuan Cao^234^, Aiguo Chen^232,235^, Huafu Chen^236^, Jie Chen^237^, Xujun Duan^236^, Xue-Ru Fan^238^, Peng Gao^238^, Wenjing Gao^239,240^, DongZhi He^239^, Feiyong Jia^229^, Haoxiang Jiang^241^, Xi Jiang^229^, Jin Jing^242^, Lei Li^236^, Shijun Li^233^, Tingyu Li^237^, Xiuhong Li^243^, Lizi Lin^242^, Yingqiang Liu^236^, Zhimei Liu^244^, Fanchao Meng^245,246^, Litong Ni^233^, Ning Pan^247^, Qi Qi^233^, Bin Qin^230^, Xiaolong Shan^236^, Xiaojing Shou^238,245,248^, Jia Wang^234^, Longlun Wang^230^, Miaoyan Wang^241^, Wei Wang^249^, Xin Wang^247^, Lijie Wu^234^, Dandan Xu^241^, Yin Xu^237^, Yang Xue^229^, Ting Yang^237^, Xuntao Yin^239^, Rong Zhang^245,248^, Yun Zhang^230^, Xi-Nian Zuo^238^

### CABIC Affiliations

^229^Department of developmental and behavioral pediatrics, Childrens Medical Center, The First Hospital of Jilin University, Jilin university, Changchun 130021, China.

^230^Department of Radiology, Children’s Hospital of Chongqing Medical University, Chongqing, 400042, China

^231^College of Physical Education, Yangzhou University, Yangzhou 225127, China

^232^School of Sport and Brain Health, Nanjing Sport Institute, Nanjing 210014, China

^233^Department of Radiology, First Medical Center, Chinese PLA General Hospital, Beijing, 100853, China

^234^Department of Childrens and Adolescent Health, Public Health College of Harbin Medical University, Harbin, 150086, China

^235^Nanjing Sport Institute, Nanjing 210014, China

^236^The Clinical Hospital of Chengdu Brain Science Institute, School of Life Science and Technology, University of Electronic Science and Technology of China, Chengdu, 610054, China.

^237^Children’s Nutrition Research Center, Ministry of Education Key Laboratory of Child Development and Disorders, National Clinical Research Center for Child Health and Disorders, China International Science and Technology Cooperation Base of Child Development and Critical Disorders, Children’s Hospital of Chongqing Medical University, Chongqing, 400042, China

^238^State Key Laboratory of Cognitive Neuroscience and Learning, Developmental Population Neuroscience Research Center at IDG/McGovern Institute for Brain Research, Beijing Normal University, Beijing, 100875, China

^239^Department of Radiology, Guangzhou Women and Childrens Medical Center, Guangzhou Medical University, Guangzhou, 511400, China

^240^School of Public Health, Shenzhen, Sun Yat-sen University, 66 Gongchang Road, Guangming District 518107, Shenzhen, China

^241^Department of Radiology, Affiliated Children’s Hospital of Jiangnan University, Wuxi, 214000, China

^242^Department of Maternal and Child Health, Joint International Research Laboratory of Environment and Health, Ministry of Education, Guangdong Provincial Engineering Technology Research Center of Environmental Pollution and Health Risk Assessment, School of Public Health, Sun Yat-sen University, Guangzhou 510080, China

^243^Department of Maternal and Child Health and Aging Health, School of Public Health, Shenzhen, Sun Yat-sen University, 66 Gongchang Road, Guangming District, 518107 Shenzhen, China

^244^Department of Rehabilitation Medicine, Affiliated Hospital of Yangzhou University, Yangzhou, 225001, China

^245^Department of Neurobiology, School of Basic Medical Sciences, Peking University Health Science Center, Beijing 100191, China

^246^The National Clinical Research Center for Mental Disorders & Beijing Key Laboratory of Mental Disorders, Beijing Anding Hospital, Capital Medical University, Beijing, China

^247^Key Laboratory of Brain, Cognition and Education Sciences, Ministry of Education, Institute for Brain Research and Rehabilitation, Guangdong Key Laboratory of Mental Health and Cognitive Science, South China Normal University, Guangzhou 510630, China

^248^Key Laboratory for Neuroscience, Ministry of Education / National Health Commission, Peking University, Beijing 100191, China

^249^Department of Radiology, Affiliated Hospital of Yangzhou University, Yangzhou, 225001, China

## SUPPLEMENTARY INFORMATION

### Supplementary Analysis

#### Brain Growth Charts and Age-Related Changes in OoS Scores for ASD Subgroup

To investigate potential developmental trajectories in ASD subgroups, here we illustrated the normative growth trajectories for volumes of the bilaterally averaged cortical regions with ABIDE dataset. The analysis focused on three brain regionstransverse temporal, inferior temporal, and isthmus cingulatethat exhibited reproducible correlations across both datasets. The subgroup-specific brain charts (Figure S1e) were upgraded from the LBCC charts and adjusted by leveraging all the samples from the ABIDE. Then, we applied Generalized Additive Mixed Models (GAMM) using the *gamm4* R package to explore the relationship between OoS scores and age within subgroup H across the CABIC and ABIDE datasets.

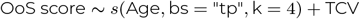

Although OoS scores are adjusted for age-related effects, examining residual variability allows us to gain additional insights into brain-cognition relationships within specific subgroups.

In the transverse temporal region (Figure S1f, left), CABIC presented a non-significant positive trend with age (*R*^2^ = 0.07, *p* = 0.56, *F* = 0.57), while ABIDE displayed a marginally significant trend (*R*^2^ = 0.06, *p* = 0.04, *F* = 4.10). For the inferior temporal region (Figure S1f, middle), CABIC showed a moderate but non-significant trend (*R*^2^ = 0.20, *p* = 0.22, *F* = 1.49), while ABIDE exhibited a flat, non-significant pattern (*R*^2^ = 0.07, *p* = 0.46, *F* = 0.56). The most robust findings were observed in the isthmus cingulate (Figure S1f, right). CABIC demonstrated a significant positive association between OoS scores and age (*R*^2^ = 0.13, *p <* 0.001, *F* = 72.69), suggesting meaningful developmental changes in this region. ABIDE also showed a positive trend, albeit with a smaller effect size (*R*^2^ = 0.12, *p* = 0.02, *F* = 5.51).

#### Spectral Clustering Independently on CABIC Cohort

To evaluate the differences between applying the ABIDE-classifier on CABIC and independently clustering analysis on CABIC, we replicated the identical analytical process directly on ASD participants from CABIC cohort. There were two distinct clusters (ASD subgroups) with the highest silhouette coefficient among cluster solutions ranging from two to ten (Figure S5a). The optimized SVM model also used a polynomial kernel with a regularization parameter *C* = 0.1 and a kernel coefficient *γ* = 0.1. The model achieved a high classification accuracy (0.97, *p <* 0.001) and a F1 score (0.97, *p <* 0.001) in five cross-validation folds. As the high cluster indices consistency (95.97%) across both methods, the brain morphology and cognitive behavior have the similar characteristics as the main analysis (Figure S5 b∼f, i∼m). Participants in subgroup L were significantly older than those in subgroup H (*p <* 0.001,). No significant differences were observed in other comparisons between two subgroups.

Significant correlations were found within two subgroups. For subgroup H, the same correlations were found as the main analysis (Figure S5g). The transverse temporal gyrus showed positive correlations with ADOS Total (*r*(201) = 0.16, *p* = 0.03) and Social Affect (*r*(193) = 0.21, *p* = 0.00) score. The inferior temporal gyrus was positively correlated with RRB (*r*(191) = 0.17, *p* = 0.02). The volumetric centile of the isthmus cingulate is correlated with SRS Autistic Mannerisms (*r*(199) = 0.15, *p* = 0.03). For subgroup L (Figure S5h), the volumetric centile of pars triangularis was positive correlations with ADOS RRB (*r*(72) = 0.24, *p* = 0.04). As for subgroup L from ABIDE cohort, *r*(86) = 0.23, *p* = 0.02.

## Supplementary Figures

**Figure S1.**
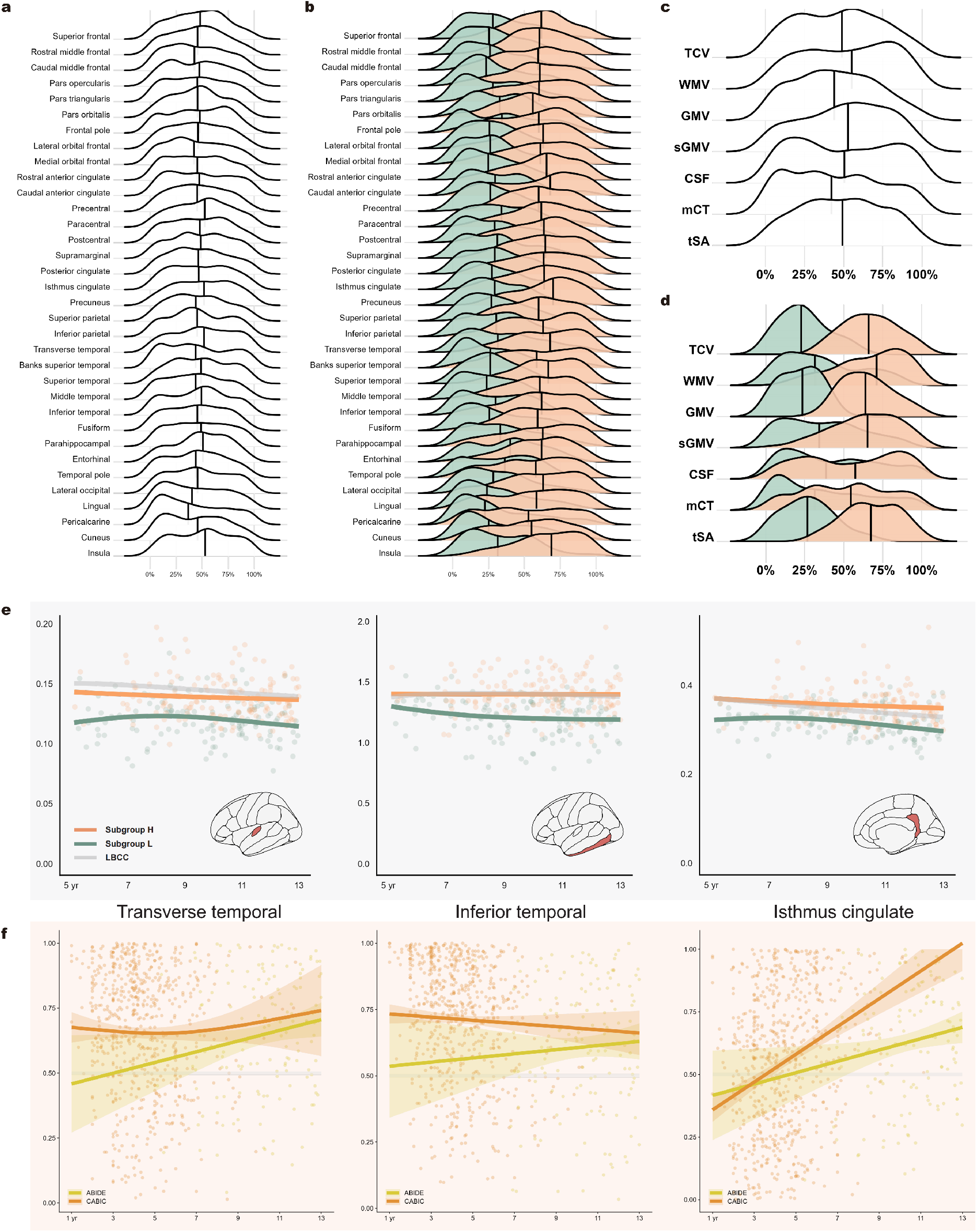
**a**, Distributions of ABIDE Out-of-Sample (OoS) scores across 34 cortical regions before clustering. **b**, Distributions of ABIDE OoS scores across 34 cortical regions after clustering. **c**, Distributions of ABIDE OoS scores across global measures before clustering. **d**, Distributions of OoS scores across global measures after clustering. **e**, ABIDE Subgroup-specific brain charts of cortical regional volumes of transverse temporal (left), inferior temporal (middle), and isthmus cingulate (right). **f**, The developmental effects of OoS scores in individuals from subgroup H were modeled using GAMM across the ABIDE and CABIC datasets. In plots **b, d**, and **e**, orange for subgroup H and green for subgroup L. TCV, total cortical volume; WMV, total white matter volume; GMV, total cortical gray matter volume; sGMV, total subcortical gray matter volume; CSF, total ventricular cerebrospinal fluid volume; mCT, mean cortical thickness; tSA, total surface area.

**Figure S2.**
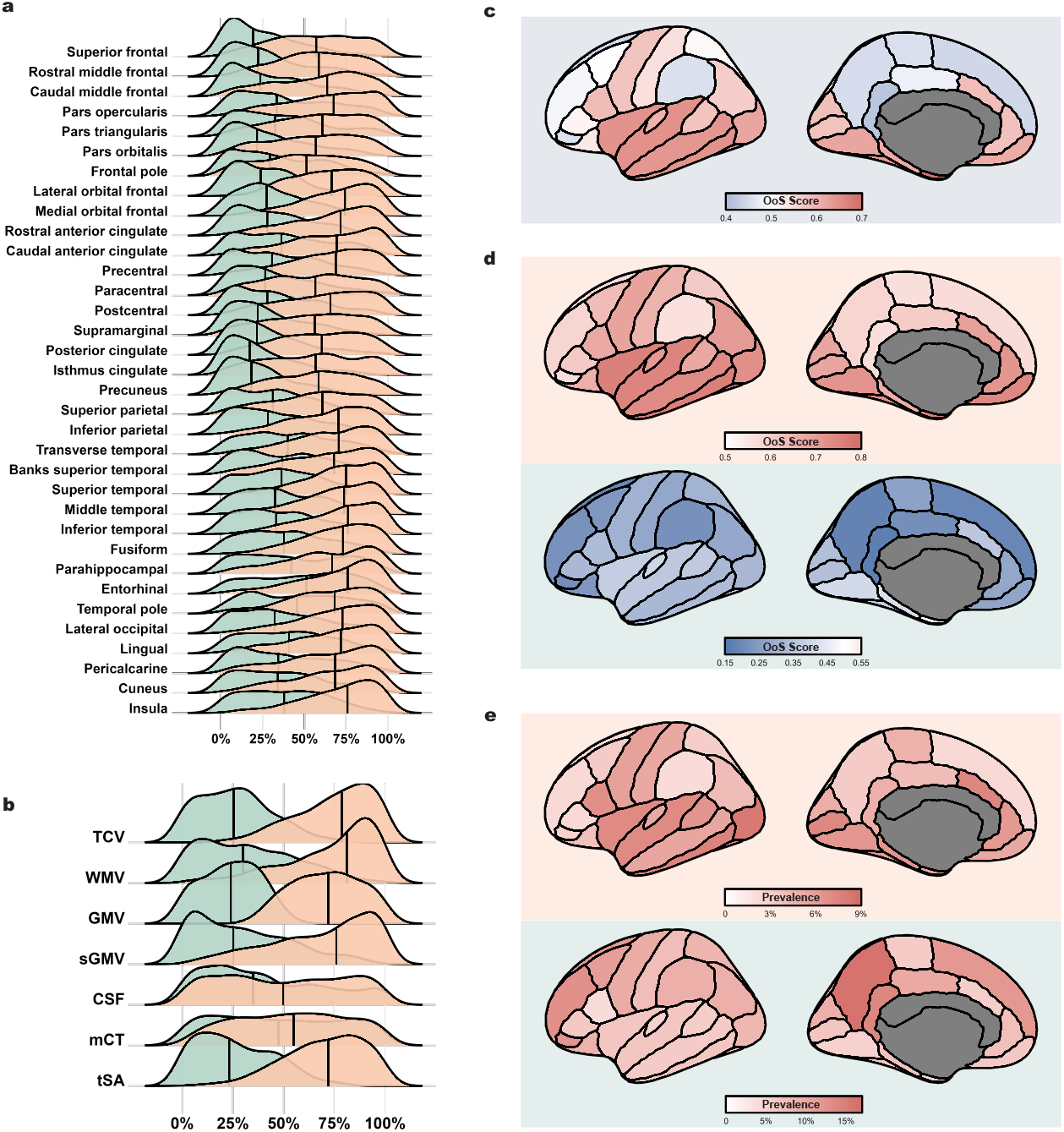
Brain morphology of two CABIC subgroups. **a**, Distributions of Out-of-Sample (OoS) scores across 34 cortical regions after clustering. **b**, Distributions of OoS scores across global measures after clustering. TCV, total cortical volume; WMV, total white matter volume; GMV, total cortical gray matter volume; sGMV, total subcortical gray matter volume; CSF, total ventricular cerebrospinal fluid volume; mCT, mean cortical thickness; tSA, total surface area. **c**, Median OoS scores of 34 before cortical regions before clustering. **d**, Median OoS scores of ASD subgroup H (top) and L (bottom). **e**, Prevalence maps depicting the proportion of participants with extreme (top, 97.5% for subgroup H; bottom, 2.5% for subgroup L) structural anomalies. Note, the left hemispheres are plotted in **c**, and **d** just for visualization purposes.

**Figure S3.**
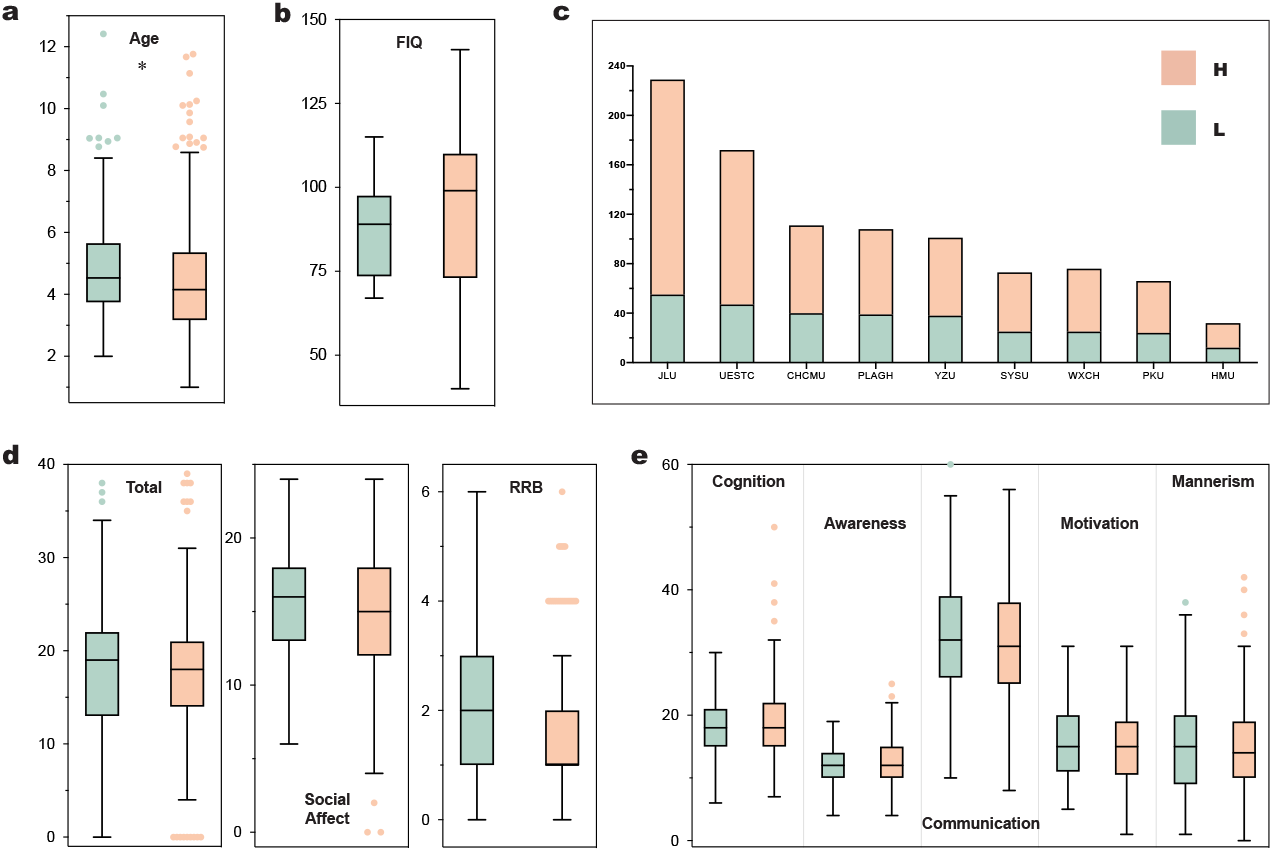
Differences between two CABIC subgroups. Distribution of participant ages (**a**); FIQ scales (**b**); data collection site (**c**); ADOS (**d**); and SRS (**e**) across the two subgroups of CABIC. Only significant difference of age (*p <* 0.001) between subgroups. For plots **a** to **e**, the center line shows the median; the box limits represent the 25th and 75th percentiles; the whiskers show the minimum and maximum values; and the dots represent potential outliers.

**Figure S4.**
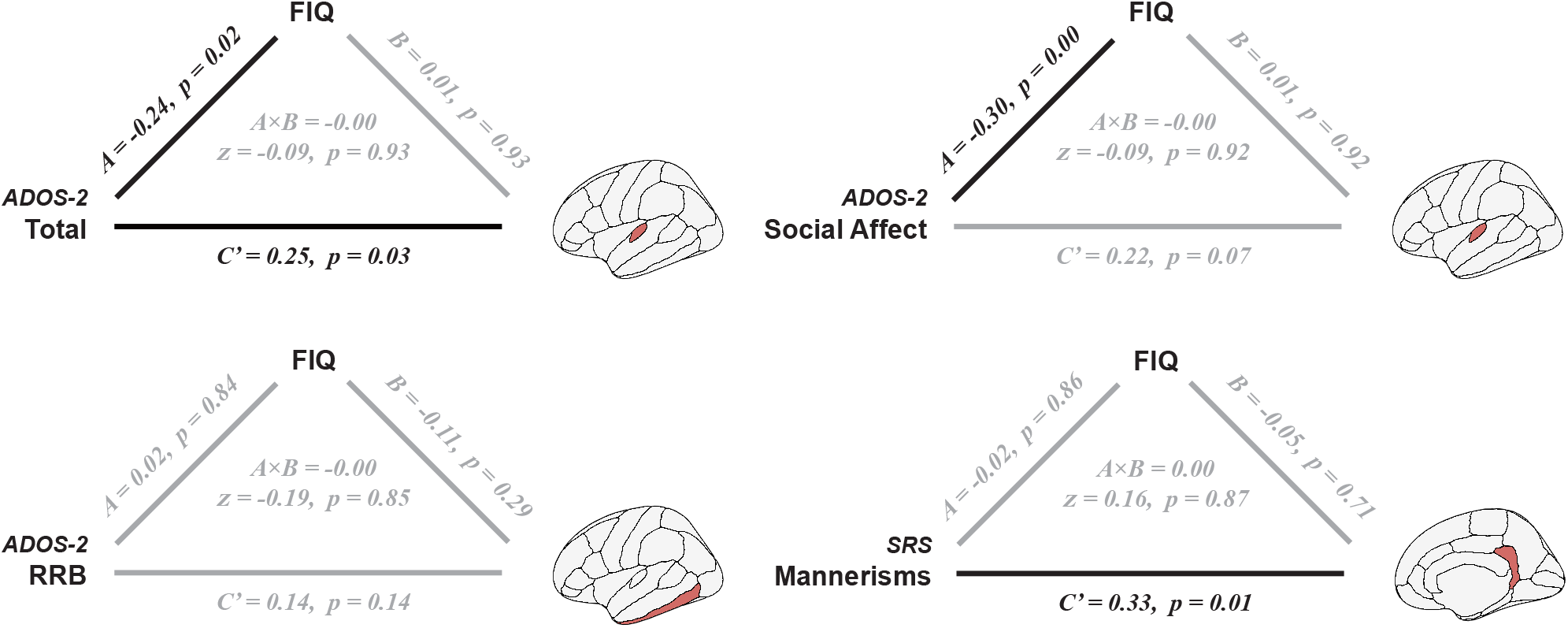
Mediation models (for significant brain-behavior associations identified in ABIDE subgroup H) to examine the cognitive behaviors’ effect on brain morphology. Black solid arrows represent significant effects, while gray arrows indicate non-significant ones. Top: transverse temporal; bottom left: inferior temporal; bottom right: isthmus cingulate.

**Figure S5.**
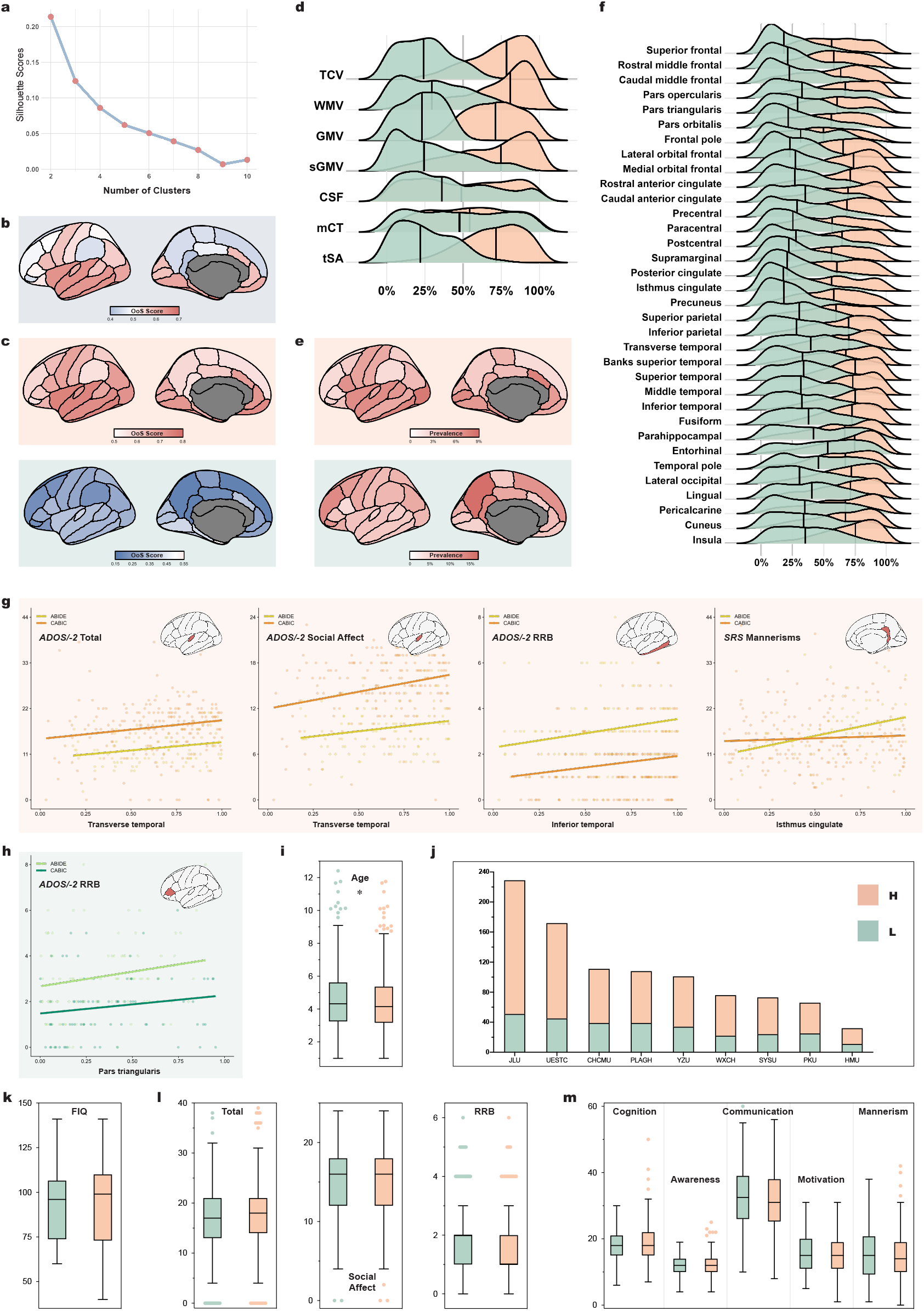
Independent spectral clustering analysis of CABIC applying the same analytical pipeline used for ABIDE. **a**, Silhouette scores across different clustering solutions (2 to 10 clusters). The highest silhouette score indicates two distinct subgroups. **b**, Median OoS scores of 34 cortical regions before clustering. **c**, Median OoS scores of ASD subgroup H (top) and L (bottom). **d**, OoS scores across global measures for subgroup H (orange) and L (green). TCV, total cortical volume; WMV, total white matter volume; GMV, total cortical gray matter volume; sGMV, total subcortical gray matter volume; CSF, total ventricular cerebrospinal fluid volume; mCT, mean cortical thickness; tSA, total surface area. **e**, Prevalence maps depicting the proportion of participants with extreme (2.5% for subgroup L, 97.5% for subgroup H) structural anomalies. **f**, OoS scores across 34 cortical regions for subgroup H (orange) and L (green). Reproducible correlations (**g**, subgroup H; **h**, subgroup L) between brain region volumes, measured as Out-of-Sample (OoS) scores, and clinical measures across ABIDE and CABIC datasets. Distribution of participant ages (**i**); data collection site (**j**); FIQ (**k**); ADOS (**l**); and SRS (**m**) across the two subgroups, with subgroup L participants being significantly older than those in subgroup H (*p <* 0.001). Note, the left hemispheres are plotted in **b, c**, and **e** just for visualization purposes. For plots **i, k, l**, and **m**, the center line shows the median; the box limits represent the 25th and 75th percentiles; the whiskers show the minimum and maximum values; and the dots represent potential outliers.

**Figure S6.**
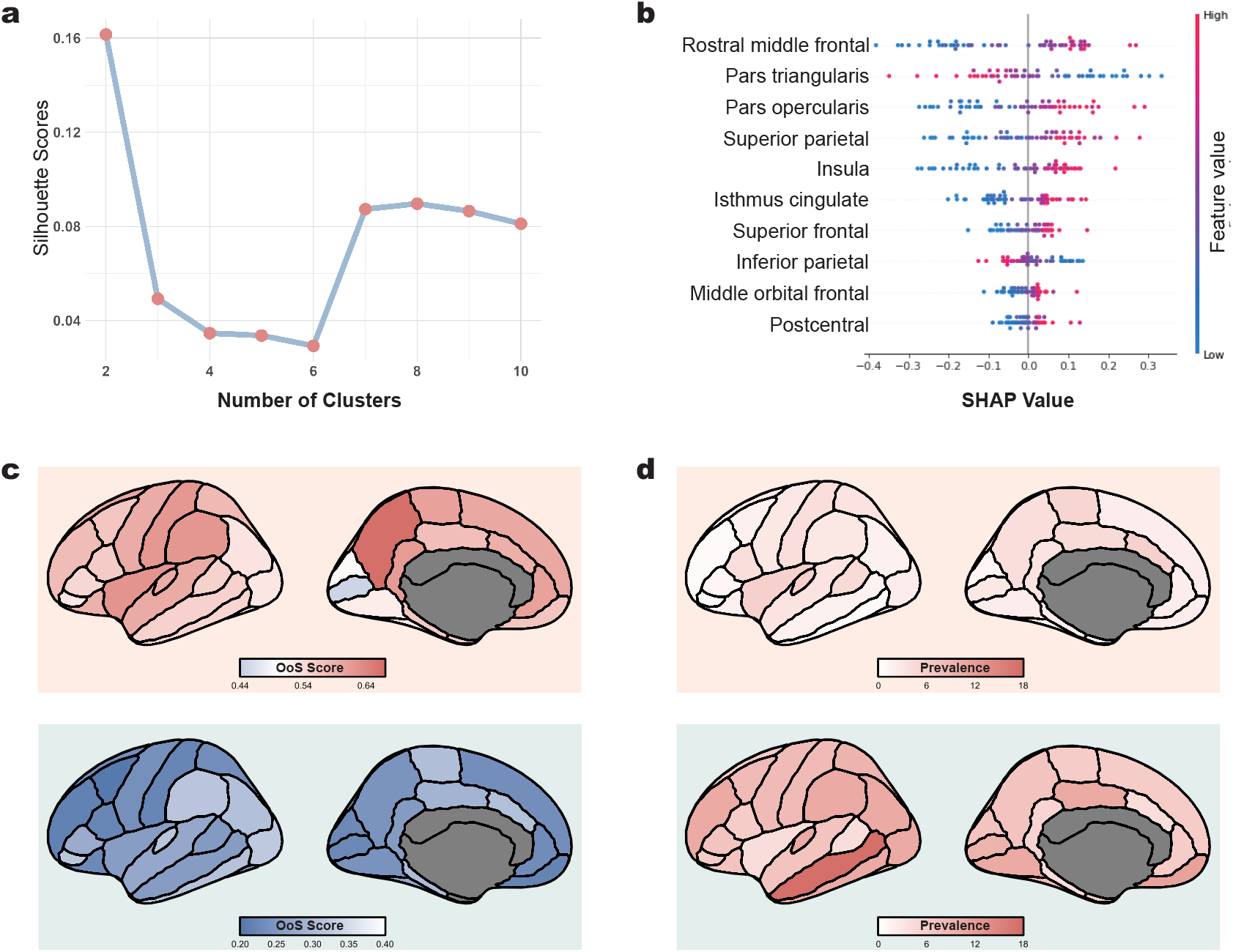
Results of GMM clustering analysis on ABIDE cohort. **a**, Silhouette scores across different clustering solutions (2 to 10 clusters). The highest silhouette score indicates two distinct subgroups. **b**, SHAP summary plot displaying the top 10 brain regions with the highest contributions to the SVM model’s predictions. **c**, Median OoS scores of ASD subgroup H (top) and L (bottom). **e**, Prevalence maps depicting the proportion of participants with extreme (top, 97.5% for subgroup H; bottom, 2.5% for subgroup L) structural anomalies. Note, the left hemispheres are plotted in **c**, and **d** just for visualization purposes.

## Supplementary Tables

**Table S1.** Statistical comparison between two clustering ASD groups for different measures.

| Phenotype |  | Group(N) | Median | Mean | SD | Shapiro-Wilk Test | Rank-Sum Test | Two-sample *t*-Test | Cohen's *d* |
| --- | --- | --- | --- | --- | --- | --- | --- | --- | --- |
| **Age** |  | H (148) | 10.51 | 10.25 | 1.87 | 0.00 | 0.02 |  | -0.30 |
|  |  | L(126) | 9.94 | 9.65 | 2.09 | 0.00 |  |  |  |
| **IQ** | PIQ | H (115) | 109 | 109.43 | 18.05 | 0.77 |  | 0.11 | -0.21 |
|  |  | L(111) | 104 | 105.76 | 16.21 | 0.19 |  |  |  |
|  | VIQ | H (116) | 110 | 109.51 | 17.23 | 0.70 |  | 0.16 | -0.19 |
|  |  | L(106) | 106 | 106.29 | 16.94 | 0.45 |  |  |  |
|  | FIQ | H (139) | 110 | 108.66 | 18.39 | 0.26 |  | 0.09 | -0.21 |
|  |  | L(117) | 104 | 105.02 | 15.83 | 0.28 |  |  |  |
| **ADOS-2** | RRB | H (108) | 3 | 3.06 | 1.84 | 0.00 | 0.94 |  | 0.02 |
|  |  | L(100) | 3 | 3.09 | 1.85 | 0.00 |  |  |  |
|  | Social Affect | H (107) | 9 | 9.36 | 3.44 | 0.00 | 0.11 |  | -0.18 |
|  |  | L(100) | 8 | 8.73 | 3.76 | 0.00 |  |  |  |
|  | Total | H (108) | 12 | 11.62 | 4.75 | 0.00 | 0.15 |  | -0.14 |
|  |  | L(100) | 11 | 11.84 | 4.75 | 0.00 |  |  |  |
|  | Severity | H (107) | 7 | 7.16 | 1.78 | 0.00 | 0.18 |  | -0.23 |
|  |  | L(100) | 7 | 6.71 | 2.14 | 0.00 |  |  |  |
| **ADI-R** | RRB | H (120) | 5 | 5.56 | 2.66 | 0.01 | 0.14 |  | 0.16 |
|  |  | L(103) | 6 | 5.96 | 2.38 | 0.02 |  |  |  |
|  | Nonverbal | H (120) | 8 | 8.12 | 2.96 | 0.15 | 0.14 |  | 0.29 |
|  |  | L(28) | 9.5 | 9.07 | 3.40 | 0.00 |  |  |  |
|  | Verbal | H (120) | 15 | 15.35 | 4.33 | 0.01 | 0.44 |  | 0.09 |
|  |  | L(103) | 16 | 15.75 | 4.35 | 0.07 |  |  |  |
|  | Social | H (119) | 19 | 18.84 | 5.52 | 0.04 | 0.77 |  | 0.04 |
|  |  | L(103) | 19 | 19.07 | 5.68 | 0.07 |  |  |  |
| **SRS** | Autistic Mannersims | H (67) | 16 | 16.49 | 6.96 | 0.53 | 0.15 |  | 0.22 |
|  |  | L(69) | 19 | 18.10 | 7.54 | 0.04 |  |  |  |
|  | Motivation | H (67) | 13 | 13.73 | 6.38 | 0.18 |  | 0.68 | 0.07 |
|  |  | L(69) | 14 | 14.17 | 6.38 | 0.52 |  |  |  |
|  | Communication | H (67) | 29 | 30.43 | 10.48 | 0.89 |  | 0.93 | -0.01 |
|  |  | L(69) | 31 | 30.28 | 11.67 | 0.38 |  |  |  |
|  | Cognition | H (67) | 17 | 16.57 | 6.60 | 0.74 |  | 0.39 | 0.15 |
|  |  | L(69) | 17 | 16.57 | 6.60 | 0.42 |  |  |  |
|  | Awareness | H (67) | 12 | 12.25 | 3.84 | 0.12 |  | 0.62 | -0.09 |
|  |  | L(69) | 11 | 11.93 | 3.73 | 0.39 |  |  |  |

**Table S2.**
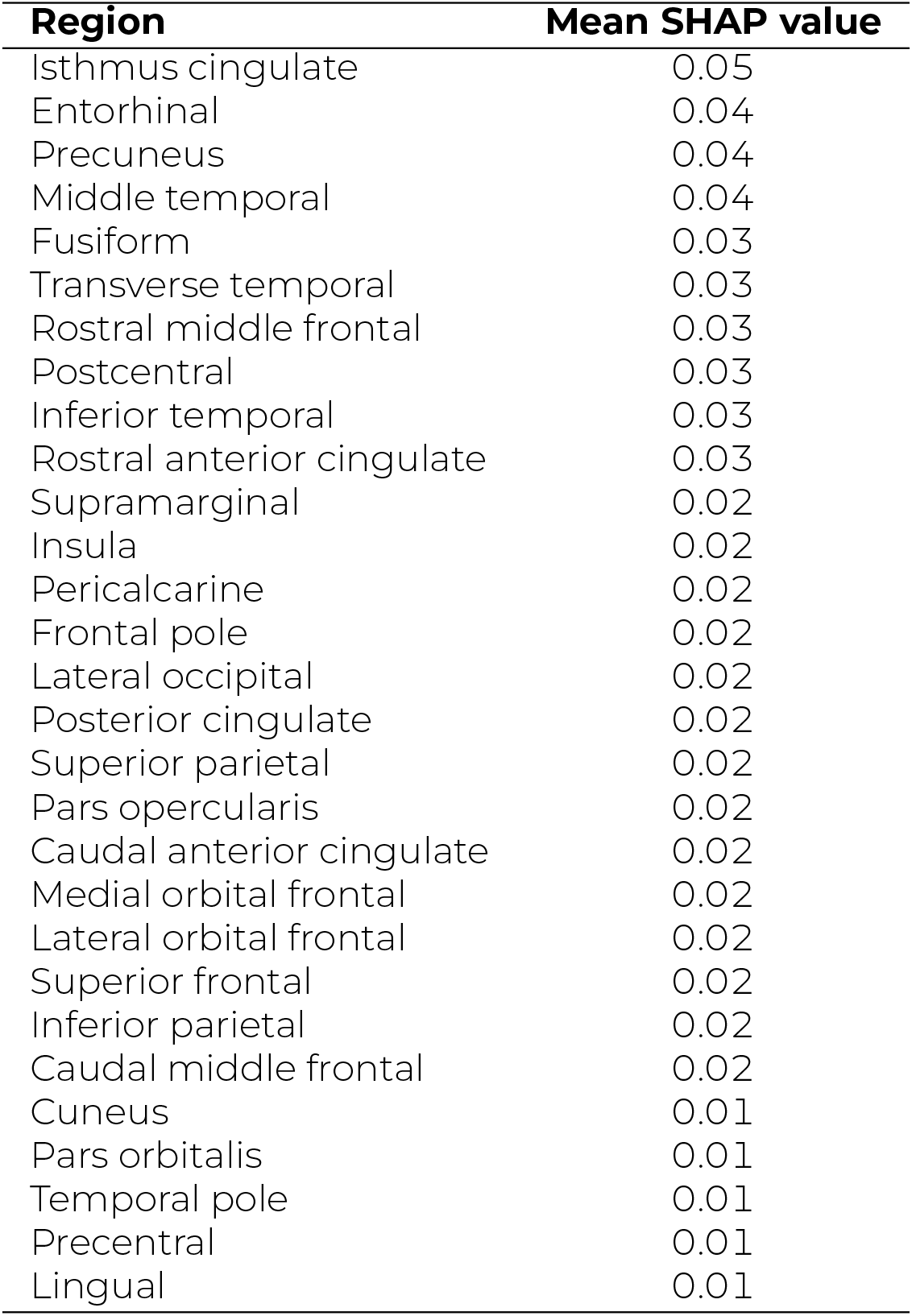
Mean SHAP values for all 29 selected regions.

## Supplementary Materials

ABIDE_ClusterID.csv

CABIC_ClusterID.csv

## Footnotes

1 AbP: Prevalence of abnormalities

